# Biases of Principal Component Analysis (PCA) in Physical Anthropology Studies Require a Reevaluation of Evolutionary Insights

**DOI:** 10.1101/2023.12.06.570437

**Authors:** Nima Mohseni, Eran Elhaik

## Abstract

Evolutionary biologists, primarily palaeoanthropologists, anatomists and ontogenists, employ modern geometric morphometrics to quantitatively analyse physical forms (e.g., skull morphology) and explore relationships, variations, and differences between samples and taxa using landmark coordinates. The standard approach comprises two steps: Generalised Procrustes Analysis (GPA) followed by Principal Component Analysis (PCA). PCA projects the superimposed data produced by GPA onto a set of uncorrelated variables, which can be visualised on scatterplots and used to draw phenetic, evolutionary, and ontogenetic conclusions. Recently, the use of PCA in genetic studies has been challenged. Due to PCA’s central role in morphometrics, we sought to evaluate the standard approach and claims based on PCA outcomes. To test PCA’s accuracy, robustness, and reproducibility using benchmark data of the crania of five papionin genera, we developed MORPHIX, a Python package for processing superimposed landmark data with classifier and outlier detection methods, which can be further visualised using various plots. Throughout this manuscript, we address the recent and contentious use of PCA in physical anthropology and phylogenetic inference, such as the case of *Homo Nesher Ramla*, an archaic hominin with a questionable taxonomy. We found that PCA outcomes are artefacts of the input data and are neither reliable, robust, nor reproducible as field members may assume. We also found that supervised machine learning classifiers are more accurate both for classification and detecting new taxa. Our findings raise concerns about PCA-based findings applied in 18,400 to 35,200 Physical anthropology studies. Our work can be used to evaluate prior and novel claims concerning the origins and relatedness of inter- and intra-species and improve phylogenetic and taxonomic reconstructions.

## Introduction

Morphometrics is one of the oldest approaches to biological investigations. It addresses various questions ranging from how shapes vary over time (evolution, growth, and development) to how they vary inter- and intra-species. It has been pursued by graphical and computational means, based on millennia-old geometric principles, since the European Renaissance (1, 2). Even though the quantitative analysis of morphological data has been long pursued (3, 4), it was not until the mid-20^th^ century that the mathematical foundations of forms matured. In the mid-1990s (5, 6, 7, 8), relevant graphic and statistical tools were made available, and modern geometric morphometrics became an essential biological tool dominating shape studies.

A major contribution to the field was made by Sokal and Sneath’s Principles of Numerical Taxonomy (9) book, which challenged traditional taxonomic theory as inherently circular and introduced quantitative methods to address questions of classification (see also review by Sneath (10)). Hull (11) claimed that evolutionary reasoning practiced in taxonomy is not inherently circular but rather unwarranted. He argued that such criticism was based on misunderstandings of the logic of hypothesising, which he attributed to an unrealistic desire for a mistake-proof science. He contended that scientific hypotheses should begin with insufficient evidence and be refined iteratively as new evidence emerges. However, some taxonomists preferred a more rigid, hierarchical approach to avoid the appearance of error. As a result of these and other criticisms, traditional taxonomy declined in favour of cladistics and molecular systematics, which provided more accurate and evolutionarily informed classifications.

Today, palaeontology and palaeoanthropology grapple with methodological challenges that compromise the stability of their conclusions. These issues stem from various factors, including biologists’ apprehensions towards advanced mathematics, the pressure to publish for career advancement (12), the pursuit of high-profile journal covers, and the prestige associated with naming new species. As a result, these fields often resemble a branch of biology where the latest discoveries or new analytical techniques frequently overturn previous findings. This lack of cumulative knowledge necessitates a more rigorous approach to methodology and interpretation in morphometrics to ensure that conclusions are robust and enduring.

Morphology is a critical source of information concerning an object’s shape and size and is used throughout the natural sciences, materials sciences, and engineering. Concerning organisms, morphometrics is the main technique for answering questions about form (shape and size) - relatedness in physical anatomy. It has many applications in evolutionary biology, anthropology, palaeontology/palaeoanthropology, and even forensics (13). Morphometrics is the leading technique in studying variations, differences, and evolutionary dynamics and relationships in forms. These variations are suggestive of ancestry and ontogeny. Field members assign morphometric ancestral and phenetic interpretations to gain ontogenetic insights into growth and development (14, 15, 16, 17, 18).

Determining sufficient measurements and data points for a valid morphometric analysis is older than modern geometric morphometrics (19). In geometric morphometrics, landmarks are discrete points on biological structures used to capture shape variation. Bookstein (20) categorised landmarks into three types: Type one, representing the juxtaposition of tissues such as the intersection of two sutures; Type two, denoting maxima of curvature like the deepest point in a depression or the most projecting point on a process; and Type three, which includes extremal points defined by information from other locations on the object, such as the endpoint or centroid of a curve or feature. Originally, Type three landmarks encompassed semi-landmarks, but Weber and Bookstein (21) refined this classification, identifying Type tree landmarks as those characterised by information from multiple curves and symmetry, including the intersection of two curves or the intersection of a curve and a suture, and further subdividing them into three subtypes (3a, 3b, 3c) (15). While landmarks provide crucial information about the structure’s overall shape, semi-landmarks capture fine-scale shape variation (e.g., curves or surfaces) that landmarks alone cannot adequately represent. Semi-landmarks are heavily relied upon as the source of shape information to break the continuity of regions in the specimen without clearly identifiable landmarks (22). Semi-landmarks are typically aligned based on their relative positions to landmarks, allowing for the comprehensive analysis of shape changes and deformations within complex structures (2). Unsurprisingly, the use of semi-landmarks is controversial. For instance, Bardua et al. (23) claim that high-density sliding semi-landmark approaches offer advantages compared to landmark-only studies, while Cardini (24) advises caution about potential biases and subsequent inaccuracies in high-density morphometric analyses.

Geometric morphometrics analysis uses different approaches that, while varying in detail, maintain similar conventional core elements (2, 16, 17, 25, 26). The standard approach to processing landmark data comprises two major steps: Generalised Procrustes Analysis (GPA) followed by Principal Component Analysis (PCA) (Figure 1). GPA superimposes the landmark coordinates by reducing the shape-independent variations in samples. PCA is a multivariate statistical analysis that projects the data onto a set of uncorrelated variables known as principal components (PCs), each explaining a reduced proportion of the variance in the data (13). After GPA, all landmark data are in the shape space manifold, meaning that each landmark configuration is a point in a high-dimensional space, where the dimensions represent the coordinates of the landmarks after GPA. However, this high-dimensional space is often too complex to visualise and analyse directly. Therefore, a tangent space projection with PCA is applied to move the points from the shape space manifold to a linearised space. In practice, only the variations along the first two or three PCs are discussed, and the rest are ignored (18, 22, 27, 28, 29, 30, 31). These PCs are then used to visualise the samples in colourful two-dimensional scatterplots to evaluate shape relationships. Typically, researchers expect biologically related shapes to cluster more closely than unrelated ones. Originally, these plots were simply developed as a tool to visually explore the structure of multivariate data in a low-dimensional subspace that summarises the bulk of shape variation along a few components (2, 20). However, the interpretation of the scatterplot patterns spread in terms of origins, relatedness, evolution, gene flow, speciation, and phenotypic/genotypic variation of the samples and taxa (16, 17, 18, 22, 26, 27, 28, 29, 30, 31, 32, 33, 34). It is also commonly assumed that the proximity of the samples is evidence of relatedness and shared evolutionary history (18, 27, 35), whereas their absence indicates the opposite. Moreover, the variation/covariation along some PCs is assumed to reflect specific morphological traits (13, 22, 36). Clearly, all these interpretations are subjective, and while some authors advised caution in interpreting the results (35), biological conclusions are commonly drawn nonetheless. For example, claims that one PC reflects ’population effects’ and another ’sex effects’ are common (e.g., 27, 32), although the PCs are statistical manifestations agnostic to the data (37).

**Figure 1:**
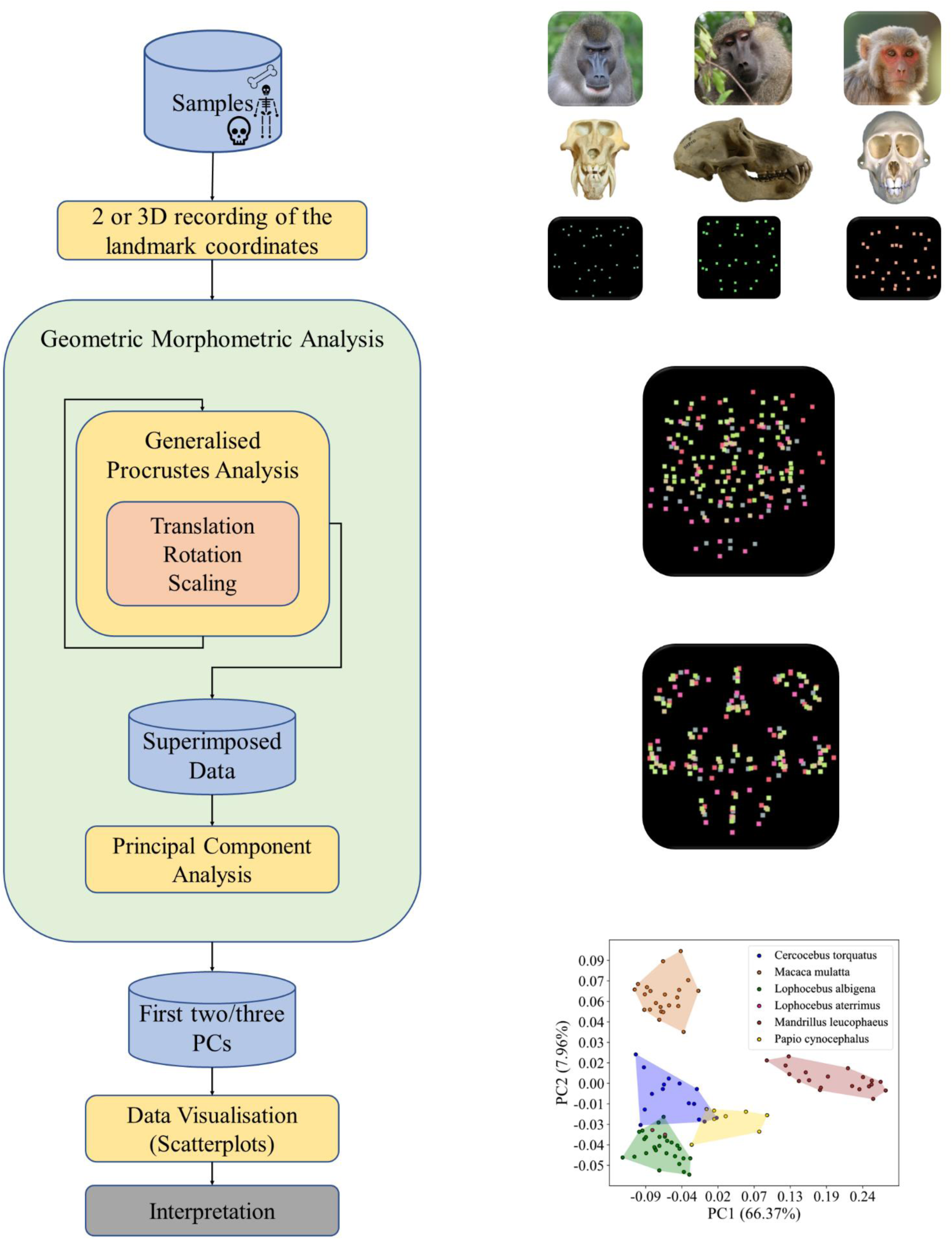
A flow chart of the conventional modern geometric morphometric analysis. The standard pipeline comprises several steps (left). First, a set of landmarks are chosen based on distinct anatomical features believed to be equivalent in some sense (e.g., biologically homologous) among all the specimens (papionins), and their cartesian coordinates are scanned and recorded. Next, the landmark data are superimposed using GPA through translation, rotation, and scaling. Next, after performing PCA, the scatterplots of the first two or three PCs are used to assess patterns of shape variation based on the distribution of the observations. The outcome of the pipeline (right) is depicted for the benchmark data. For brevity, only three taxa are shown on the top right.

Morphology-based classification, historically a cornerstone of palaeoanthropology (38), has evolved with modern geometric morphometrics. One popular application of modern geometric morphometrics is the classification of newly discovered hominin fossils within the human phylogenetic tree by visually observing the *clustering* and *proximity* of the specimens with each other in PC plots (18, 22, 27) in conjunction with other analyses. However, which specimens should be included in *clusters* and which ones should be considered *outliers* is determined subjectively. Similarly, plots produced by different PCs might yield inconsistent results, and authors may choose plots based on subjective and inconsistent criteria. In the following, we will reflect on three recent and controversial use cases of morphometrics and where PCA has been misused for phylogenetic inference. In the first case of *Nesher Ramla* (NR) fossils bones found in Israel, Hershkovitz et al. (22) compared the osteological remains with those of other hominins using PC1, PC2, and PC3 scatterplots to investigate their origins (Figures 2, 3, S1, and S17 in (22)). The plots produced conflicting results. Excluded from that study was the lower molar PC2-PC3 plot shown in Figure 2C. In this plot, NR fossils clusters with *Neanderthals*, which was inconsistent with the PC1-PC2 (Figures 2A, Figure S17 A in (22)) and PC1-PC3 (Figures 2B, Figure S17 B in (22)) plots, even though shape variation along both PC2 and PC3 was relied upon and interpreted in the text (22). Selecting a few PC dimensions and excluding the conflicting results, the authors concluded that the Nesher Ramla fossils were among the last survivors of an as-of-yet-unknown population contributing to the Neanderthals and East Asian *Homo*. A similar methodology was used by Wu et al. (2023) for the Hualongdong sample 6 (HLD 6), comprising a mandible and partial crania (18). The authors analysed the clustering, overlap, and position of HLD 6 relative to the samples in PC scatterplots (18). They reported that the overall mandibular shape and size of HLD 6 has an intermediary position among five other overlapping groups in the PC1-3 scatterplot. Moreover, in the PC1-2 scatterplot of the shape of the symphysis, HLD 6 was located in the middle of the overlap of Neanderthals and the Late Pleistocene modern humans in PC2. Along the PC1 axis, HLD 6 was located at the edge of the range of variation of the Middle Pleistocene hominins and recent modern humans (18). Based on these observations, the authors claimed that HLD 6 could belong to a new branch in the human family tree. Finally, PCA also played a part in the much-disputed case of the Dmanisi hominins (39, 40). These early Pleistocene hominins, whose fossils were recovered at Dmanisi (Georgia), have been a subject of intense study and debate within physical anthropology. Despite their small brain size and primitive skeletal architecture, the Dmanisi fossils represent Eurasia’s earliest well-dated hominin fossils, offering insights into early hominin migrations out of Africa. The taxonomic status of the Dmanisi hominins has been initially classified as *Homo erectus* or potentially represented a new species, *Homo georgicus* or else (40, 41). Lordkipanidze et al.’s (42) geometric morphometrics analyses suggested that the variation observed among the Dmanisi skulls may represent a single regional variant of Homo erectus. However, Schwartz et al. (43) raised concerns about the phylogenetic inferences based on PCA results of the geometric morphometrics analysis, noting the failure of the method to capture visually obvious differences between the Dmanisi crania and specimens commonly subsumed under *Homo erectus*.

**Figure 2:**
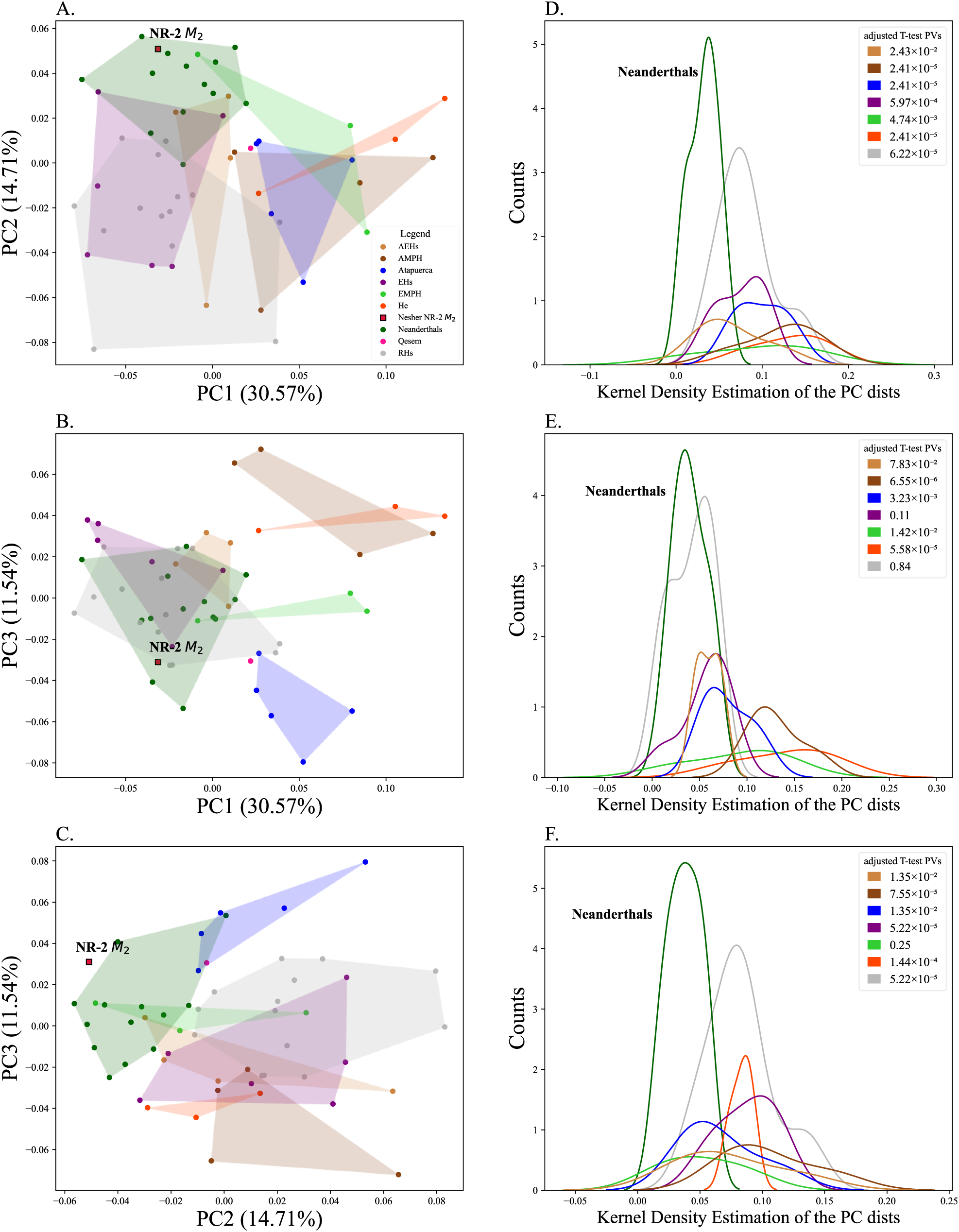
PC scatterplots of the shape space of the lower molar of NR2 and other *Hominins* (left). The analysis combines the enamel–dentine junction (EDJ) and the cemento-enamel junction (CEJ). The Kernel Density Estimation (KDE) plots the distribution of the distances of each *Homo* species from *NR* fossils in the space of the corresponding PCs (right). The legends of the right sub-figures present the adjusted (Benjamini/Hochberg) p-values of the T-test for the means of the distances of each *Hominin* population and *Neanderthals* from *NR* fossils. Legend guide: African Early *Homo sapiens* (AEHs), African *Middle Pleistocene Homo* (MPH), Early *Homo sapiens* (EHs), European *Middle Pleistocene Homo* (EMPH), *Homo erectus* (He), Recent *Homo sapiens* (RHs). For consistency, the colours of the species are similar to those of Hershkovitz et al. (22) PC plots.

Visual interpretations of the PC scatterplots are not the only role PCA plays in geometric morphometrics. Phylogenetic Principal Component Analysis (Phy-PCA) (44) and Phylogenetically Aligned Component Analysis (PACA) (45) are both used in geometric morphometrics to analyse shape variation while considering the supposed phylogenetic relationships among species. They differ in their approach to aligning landmark configurations and the role of PCA within them. Phy-PCA incorporates phylogenetic information by utilising a phylogenetic tree to model the evolutionary history of the species. This method aims to separate shape variation resulting from shared evolutionary history from other sources of variation. PCA plays a similar role in performing dimensionality reduction on the aligned landmark configurations in Phy-PCA (44). PACA takes a different approach to alignment. It uses a Procrustes superimposition method based on a phylogenetic distance matrix, aligning the landmark configurations according to the evolutionary relationships among species. PCA is then applied to the aligned configurations to extract the principal components of shape variation (45). Both analyses provide insights into the patterns and processes that shape biological form diversity while considering phylogenetic relationships, yet they are also subjected to the limitations and biases inherent in relying on PCA as part of the process.

Criticism of PCA in morphometrics is not new. Rohlf (46) already observed that distances between close neighbours are not well represented by PCA ordination. However, due to the lack of satisfactory methods for deriving clusters from similarity matrices (Sokal and Sneath 1963 (9), Pg. 252), and the premise of using a quantitative method, PCA usage continued and spread despite its stated limitations. No sooner had PCA been established as the main element of geometric morphometric analysis that even the criticism from its pioneers was dismissed (5). More recent concerns about the adequacy of PCA (47) and its extensions (5, 48) in morphometrics and phylogenetics were also published. It has been shown that between-group PCA can inflate and even create differences and patterns that do not exist (48) and that common PCA (multi-group PCA) (49) introduces systematic statistical biases in morphological trait analysis, even when the data are in very good condition (50). Furthermore, through evaluating numerous cases using population genomics data, it has been shown that projections from high to low dimensional space shown in PC scatterplots are artefacts of the input data and can produce a multitude of outcomes based on variation in the input data (51). These artefacts are not limited to the nearest neighbours, as Rohlf (108) argued, but also include distances between the major clusters in subsets of PCs, usually restricted to the first two. Despite the ongoing criticism and concerns surrounding PCA in morphometrics, a substantial lack of organised scepticism persists in the literature, highlighting the need for further critical evaluation and scrutiny of the methodologies employed in geometric morphometric analysis.

PCA’s prominent role in morphometrics analyses and, more generally, physical anthropology is inconsistent with the recent criticisms, raising concerns regarding its validity and, consequently, the value of the results reported in the literature. To assess PCA’s accuracy, robustness, and reproducibility in geometric morphometric analysis, particularly its potential biases and inconsistencies in clustering with species taxonomy for phylogenetic reconstruction, we utilised a benchmark database containing landmarks from six known species within the Old-World monkeys tribe Papionini. We altered this dataset to simulate typical characteristics of paleontological data. We found that PCA’s outcomes lack reliability, robustness, and reproducibility. We also evaluated the argument that a high explained variance could be counted as a measure of reliability (2) and found no association between high explained variance amounts and the subjectiveness of the results. If PCA of morphometric landmark data produces biased results, then landmark-based geometric morphometric studies employing PCA, conservatively estimated to range from 18,400 to 35,200 (as of July 2024) (see Methods), should be reevaluated.

As an alternative to the PCA step in geometric morphometrics analysis for species taxonomical clustering for phylogenetic reconstruction facilitation, we assessed leading supervised learning classifiers and outlier detection methods using the original and altered three-dimensional landmark datasets. We found that the nearest neighbour classifier and local outlier factor outperformed all other methods in classification and outlier detection, respectively. Concerningly, we also observed that the accuracy of linear discriminant analysis (LDA), a classifier already utilised in geometric morphometrics, was strongly influenced by the selection of the number of principal components (PCs) used as input. We developed MORPHIX, a Python package that contains the necessary tools for processing superimposed landmark data with any classifier and outlier detection method of choice, which can be further visualised using various plots. MORPHIX is freely and publicly available at https://github.com/NimaMohseni/Morphometric-Analysis and includes numerous detailed examples.

## Results

We compared the classification accuracy of PC scatterplots and eight Machine Learning (ML) tools for the complete dataset of the crania of 94 papionin adults belonging to six species (*Cercocebus torquatus, Papio cynocephalus, Lophocebus albigena*, *Lophocebus aterrimus, Macaca mulatta,* and *Mandrillus leucophaeus*) with 31 three-dimensional landmarks (the benchmark) (Figure 3). Each of the four cases of altered datasets (species removal, sample removal, landmark removal, and their combination) includes two examples. If PC-based shape inferences are reliable (16, 22, 26, 27, 29) and if PC scatterplots are valid tools for classifying samples and investigating phylogeny (17, 18, 52, 53), as members of the field assume, different species should not overlap in PC space, whereas related species who share recent evolutionary origins are expected to cluster, overlap, or exhibit short distances from each other and the distribution of samples and groups along PCs should accurately reflect relatedness in certain morphological traits (22, 27).

**Figure 3.**
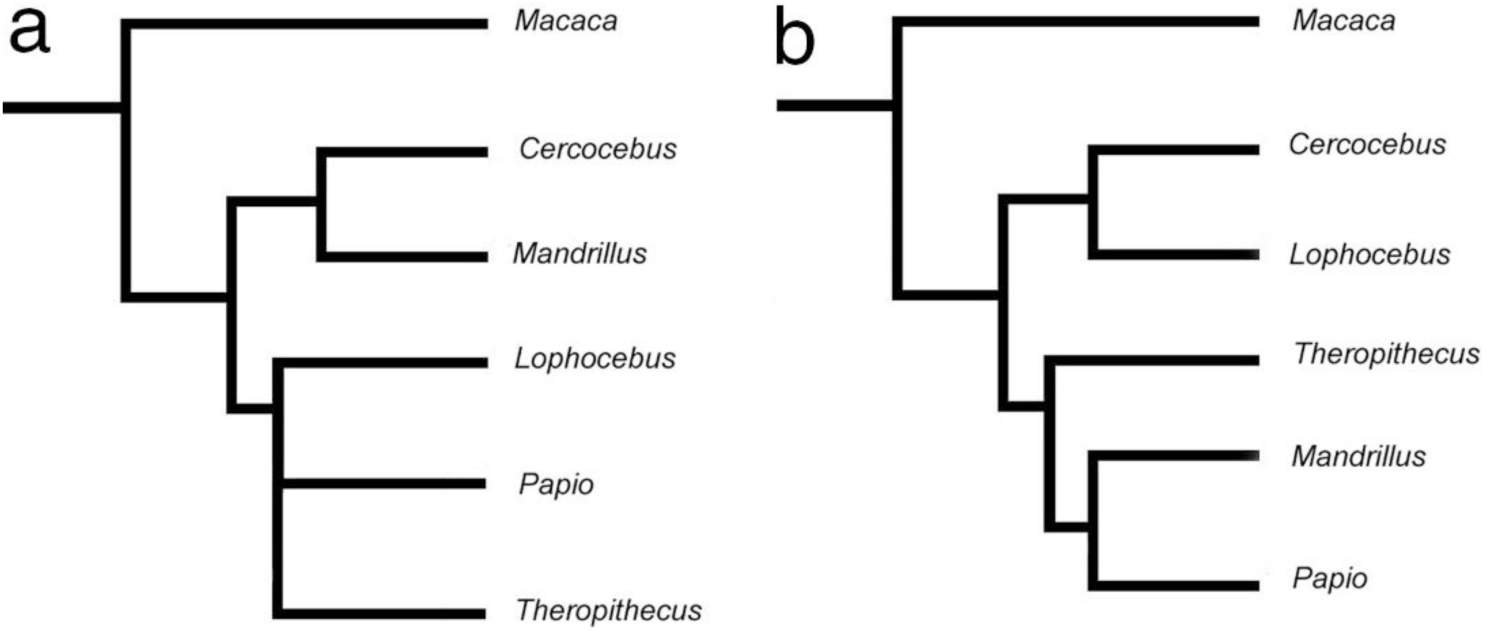
Comparing two hypotheses of papionin phylogenetic relationships. (*a*) Hypothesised phylogenetic tree of the extant Papionini from molecular (mtDNA and Y chromosome) data. (*b)* the most parsimonious tree, using *Macaca* rather than *Pan* as the outgroup. The figure was obtained from Gilbert et al. (105) (their Figure 2) under the exclusive PNAS License to Publish^1^.

### The benchmark data

First, we carried out traditional geometric morphometric analyses using GPA and PCA. Next, we reanalysed the data using alternative classifiers. Even without any alterations, the PC-based clusters are inconsistent with the taxonomical knowledge of the papionin taxa (Figure 3). Any standard (e.g., (44)) phylogenetic inference based on these PC clusters will necessarily produce an erroneous phylogeny. In all PC plots (Figures 4A-C), at least two groups (not necessarily the same ones) overlap. For example, in Figure 4A *Cercocebus torquatus (blue)* overlaps with both *Papio cynocephalus (yellow)* and *Lophocebus albigena (green)*, in Figure 4B *Macaca mulatta (orange)*, *Cercocebus torquatus (blue)* and *Lophocebus albigena (green)*, and in Figure 4C *Mandrillus leucophaeus (red)* and *Cercocebus torquatus (blue)* overlap. Consequently, and since the results can be expected to continue to change as higher dimensions are explored, it is unclear whether these overlapping samples should be treated as misclassification as there is room for other evolutionary interpretations. Based on the proximity of the samples to their neighbours, misinterpretations can range from phenotypic variation (16, 54), admixture and hybridisation (26), to speciation (17) and gene flow (22) and phenetic evolutionary trends (18). Observations can be further explained in different manners. For example, interpreting the results as gene flow might also require explaining its direction; this was the case of *NR* fossils, whom, as Hershkovits et al. claimed (22), could have belonged to a population ancestral to the *H. neanderthalensis* and East Asian *Homo,* while Rightmire argued (33) that *NR* fossils could have belonged to a group of European *H. neanderthalensis* who had migrated to the area based on the same findings. Considering the *Cercocebus torquatus (blue)* samples that fall near or within *Lophocebus albigena (green)* in Figure 4A; if they were newfound samples of unknown origins, they could be misinterpreted as being *Lophocebus albigena (green)*. Additionally, several different gene flow scenarios could also be proposed, such as the *Lophocebus albigena (green)*-proximate *Cercocebus torquatus* samples being from an as-of-yet unknown origin contributing to the *Cercocebus torquatus* (blue) and *Lophocebus albigena* (green) or being a hybrid of them.

**Figure 4:**
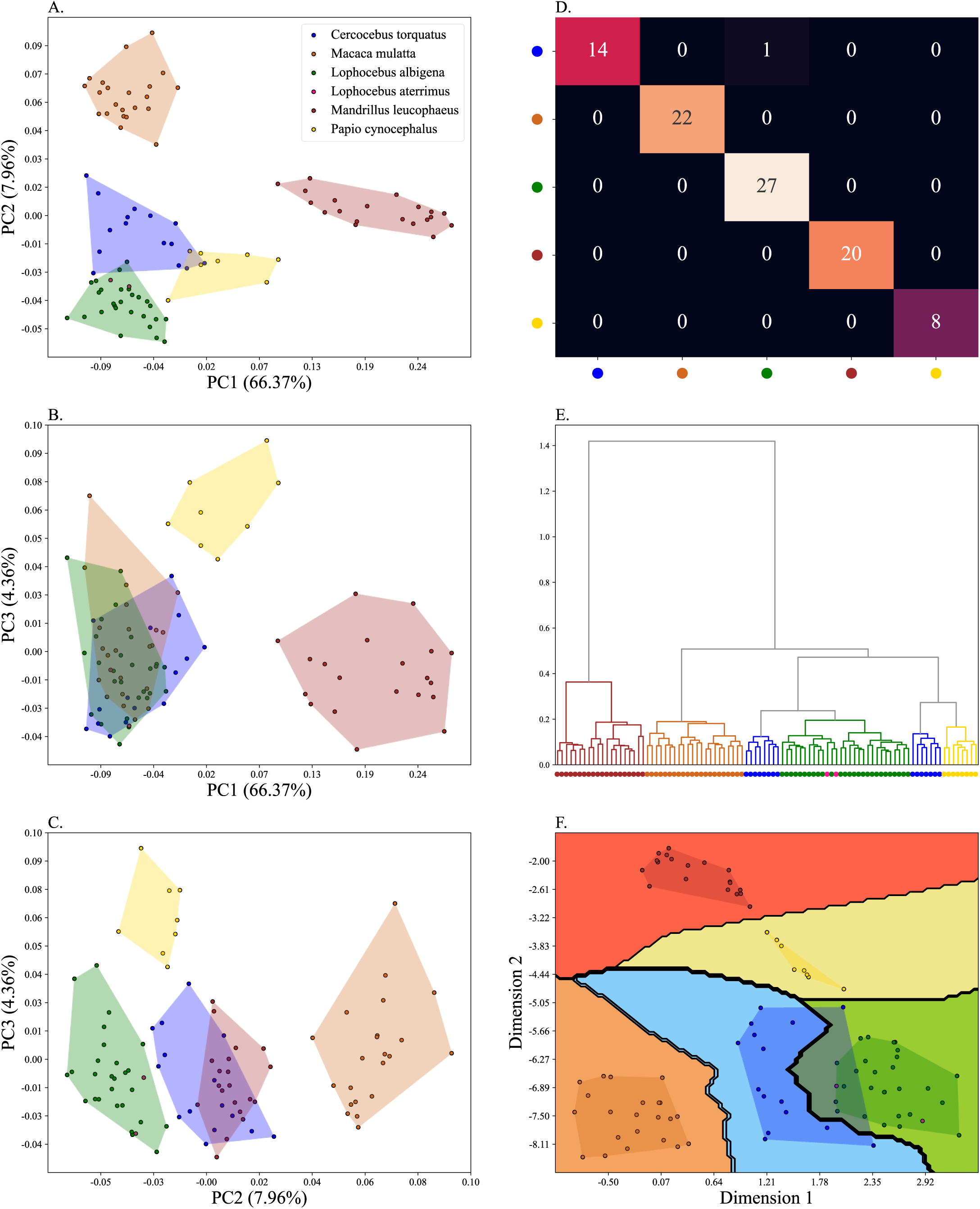
Plots depicting the papionin benchmark data. A) PC 1-2, B) PC 1-3, C) PC 2-3, D) Confusion matrix of a 2NN classifier with 5-fold CV, E) Dendrogram of Procrustes distances of the samples using agglomerative clustering, F) Decision boundary of the 2NN classifier after t-SNE embedding with 5-fold CV.

It is customary to interpret the sum of the explained variance of the PCs as a measure to evaluate the accuracy (28, 29). Despite occasional advised caution (55, 56), higher percentages of explained variance are still interpreted as reflecting higher accuracy (57, 58). We first show that this is not always the case. In Figure 4A, PCs can have the highest explained variance (Figure 4A, PC1+PC2=74%), yet the overlap of taxa *Cercocebus torquatus (blue)* and *Papio cynocephalus (yellow)* would necessarily lead to the aforementioned misinterpretations. By contrast, in Figure 4C, with the lowest explained variance (explained variance =12%), the PCs separate *Cercocebus torquatus (blue)* and *Papio cynocephalus (yellow)*. In other words, by pure chance, here, the PC plot with the lesser variance depicts morphometrical relatedness and distances more adequately and accurately than the plot, explaining shape variation more than six times.

Next, we investigated the classification *accuracy* of the alternative methods in classifying superimposed landmark data. We calculated the accuracy as the portion of the correctly classified samples to the total number of samples and *balanced accuracy* as the arithmetic mean of sensitivity and specificity, which can be generalised to multiclass classification under one versus-rest transformation (59, 60). We implemented eight ML methods that could effectively classify the samples in a five-fold Cross-Validation (CV) scheme, including Linear Discriminant Analysis (LDA), using the first two and 10 PCs of the superimposed landmarks. To assess the ability of PC scatterplots to classify samples (17, 52, 53), we defined accuracy as the ratio of the non-overlapping samples to all samples in each case. Since, in case of overlap, false negatives cannot be defined, no balanced accuracy could be calculated for PC plots. It should be emphasised that LDA was solely used as a classifier and not a dimensionality reduction method.

Three ML classifiers, including the Logistic regression (LR), Gaussian Process (GP), and Support Vector Classifier (SVC), as well as the Nearest Neighbours (NN) (Figure 4F), had balanced accuracies higher than 96%. The three classifiers from the decision tree family had lower performances. LDA had an accuracy of 93% and 98% for the first two and ten PCs, respectively. We visualised the samples and the decision boundaries of the most effective classifier using a t-distributed Stochastic Neighbour Embedding (t-SNE) (61). However, we note that the main infographic sources should be the confusion matrices (supplementary Figures S1-9) and Procrustes distances dendrograms created using agglomerative clustering (90). Table 1 presents the accuracy and balanced accuracy of each classifier.

**Table 1:** The accuracy and balanced accuracy of the eight classifiers. The accuracy was evaluated in a five-fold stratified CV on the benchmark dataset and the six alteration cases. The numbers in parentheses are the balanced accuracies.

|  | NN | LR | GP | SVC | RF | ET | XGB | LDA(2) | LDA(10) | PC1-2 | PC1-3 | PC2-3 |
| --- | --- | --- | --- | --- | --- | --- | --- | --- | --- | --- | --- | --- |
| **Benchmark data** | 0.97<br>(0.97) | 0.97<br>(0.96) | 0.98<br>(0.98) | 0.97<br>(0.97) | 0.92<br>(0.88) | 0.90<br>(0.86) | 0.92<br>(0.91) | 0.93<br>(0.90) | 0.98<br>(0.98) | 0.93<br>- | 0.42<br>- | 0.79<br>- |
| **Removing Green** | 1<br>(1) | 0.98<br>(0.96) | 1<br>(1) | 1<br>(1) | 0.96<br>(0.95) | 0.96<br>(0.95) | 0.95<br>(0.92) | 0.92<br>(0.92) | 1<br>(1) | 0.97<br>- | 0.98<br>- | 0.92<br>- |
| **Removing Orange** | 1<br>(1) | 1<br>(1) | 1<br>(1) | 1<br>(1) | 0.92<br>(0.90) | 0.94<br>(0.93) | 0.9<br>(0.88) | 0.8<br>(0.79) | 1<br>(1) | 0.62<br>- | 0.75<br>- | 0.37<br>- |
| **Green sample removal** | 0.98<br>(0.98) | 0.97<br>(0.96) | 0.98<br>(0.98) | 0.98<br>(0.98) | 0.97<br>(0.97) | 0.98<br>(0.98) | 0.95<br>(0.93) | 0.91<br>(1) | 0.88<br>(1) | 0.91 | 0.5 | 0.96 |
| **Yellow and Blue sample removal** | 0.97<br>(0.94) | 0.93<br>(0.86) | 0.97<br>(0.94) | 0.98<br>(0.97) | 0.95<br>(0.91) | 0.95<br>(0.91) | 0.90<br>(0.85) | 0.91<br>(0.88) | 1<br>(1) | 0.93 | 0.54 | 1 |
| **landmark removal (23, 24, 25, 26 and 27)** | 0.98<br>(0.98) | 0.96<br>(0.94) | 0.98<br>(0.98) | 0.97<br>(0.97) | 0.95<br>(0.93) | 0.93<br>(0.90) | 0.90<br>(0.84) | 0.91<br>(0.90) | 1<br>(1) | 0.90<br>- | 0.38<br>- | 0.78<br>- |
| **landmark removal (11, 12, 16, 17, 18, 30 and 31)** | 0.95<br>(0.95) | 0.93<br>(0.91) | 0.95<br>(0.95) | 0.94<br>(0.93) | 0.90<br>(0.88) | 0.91<br>(0.89) | 0.85<br>(0.81) | 0.93<br>(0.90) | 0.96<br>(0.96) | 0.87<br>- | 0.48<br>- | 0.76<br>- |
| **Extreme case 1** | 0.71<br>(0.57) | 0.71<br>(0.54) | 0.64<br>(0.46) | 0.67<br>(0.52) | 0.71<br>(0.54) | 0.75<br>(0.59) | 0.64<br>(0.52) | 0.71<br>(0.59) | 0.71<br>(0.63) | -<br>- | -<br>- | -<br>- |
| **Extreme case 2** | 61<br>(0.51) | 0.5<br>(0.38) | 0.58<br>(0.42) | 0.52<br>(0.41) | 0.6<br>(0.54) | 0.55<br>(0.47) | 0.58<br>(0.43) | 0.41<br>(0.34) | 0.47<br>(0.42) | -<br>- | -<br>- | -<br>- |

### The case of missing taxa

Paleontological findings often differ in quality and completeness, which poses profound challenges to morphometric analyses. Rarely are all related taxa represented; thereby, only partial datasets are inevitably used (18, 22, 27). Consequently, the extent of the bias created by this incomplete sampling is unclear. To study how PCA and alternative classifiers are affected by non-representative partial sampling, we excluded one taxon at a time from the dataset, applied the pipeline (Figure 1) and classifiers, and compared the outcome to the benchmark (Figure 4).

We present two instances where the removal of *Lophocebus albigena (green)* (Figure 5) and *Macaca mulatta (orange)* (Figure 6) resulted in profound deviations compared to the benchmark. In both cases, the distances within and among groups in the PC plots changed dramatically. Removing the *Lophocebus albigena (green)* results in three instances of taxa overlap (separated in the benchmark). *Macaca mulatta (orange)* overlaps with *Cercocebus torquatus (blue)* in Figure 5A and Figure 5B and with *Mandrillus leucophaeus (red)* in Figure 5C. In the case of the *Macaca mulatta (orange)* removal, the changes in PC plots are even more profound (Figure 6). Here *Cercocebus torquatus (blue)* and *Lophocebus albigena (green)* clusters overlap almost entirely across all the PC plots (Figures 6A -6C), whereas they had a minimal overlap in the benchmark plot (Figure 5A). Another noticeable change is the overlap of *Macaca mulatta (orange)* and *Mandrillus leucophaeus (red)*. While in the benchmark, they are completely separated by PC2 (Figures 4A and 4C), after the removal of *Lophocebus albigena (green)*, *Macaca mulatta (orange)* overlaps considerably with *Mandrillus leucophaeus (red)* (Figures 6A and 6C).

**Figure 5:**
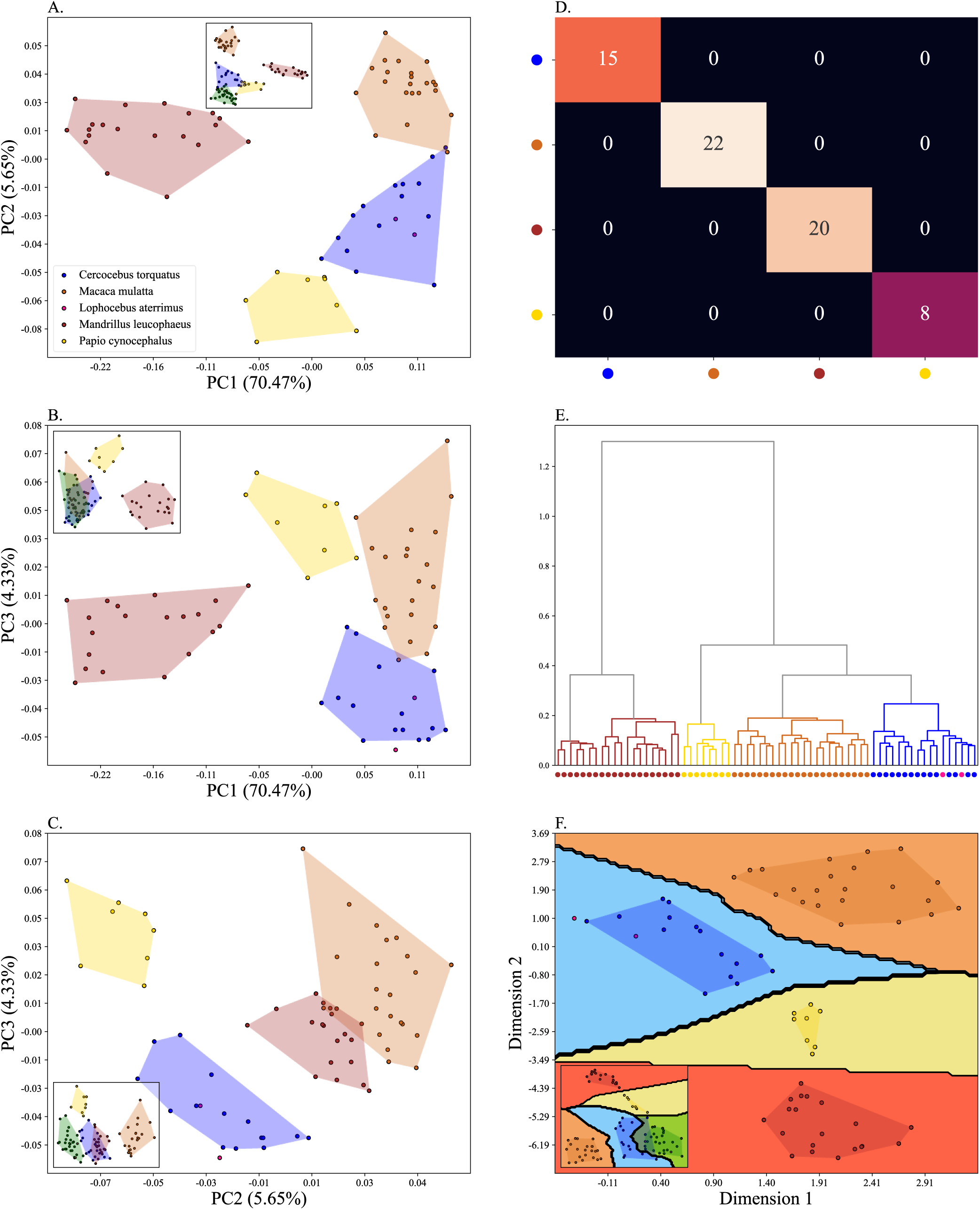
Plots depicting the papionin data after the removal of the Lophocebus albigena (green) taxon. A) PC 1-2, B) PC 1-3, C) PC 2-3, D) Confusion matrix of a 2NN classifier with 5-fold CV, E) Dendrogram of Procrustes distances of the samples using agglomerative clustering, F) Decision boundary of the 2NN classifier after t-SNE embedding with 5-fold CV. The insets in the subfigures are the corresponding benchmark PC scatterplots (Figure 4A-C).

**Figure 6:**
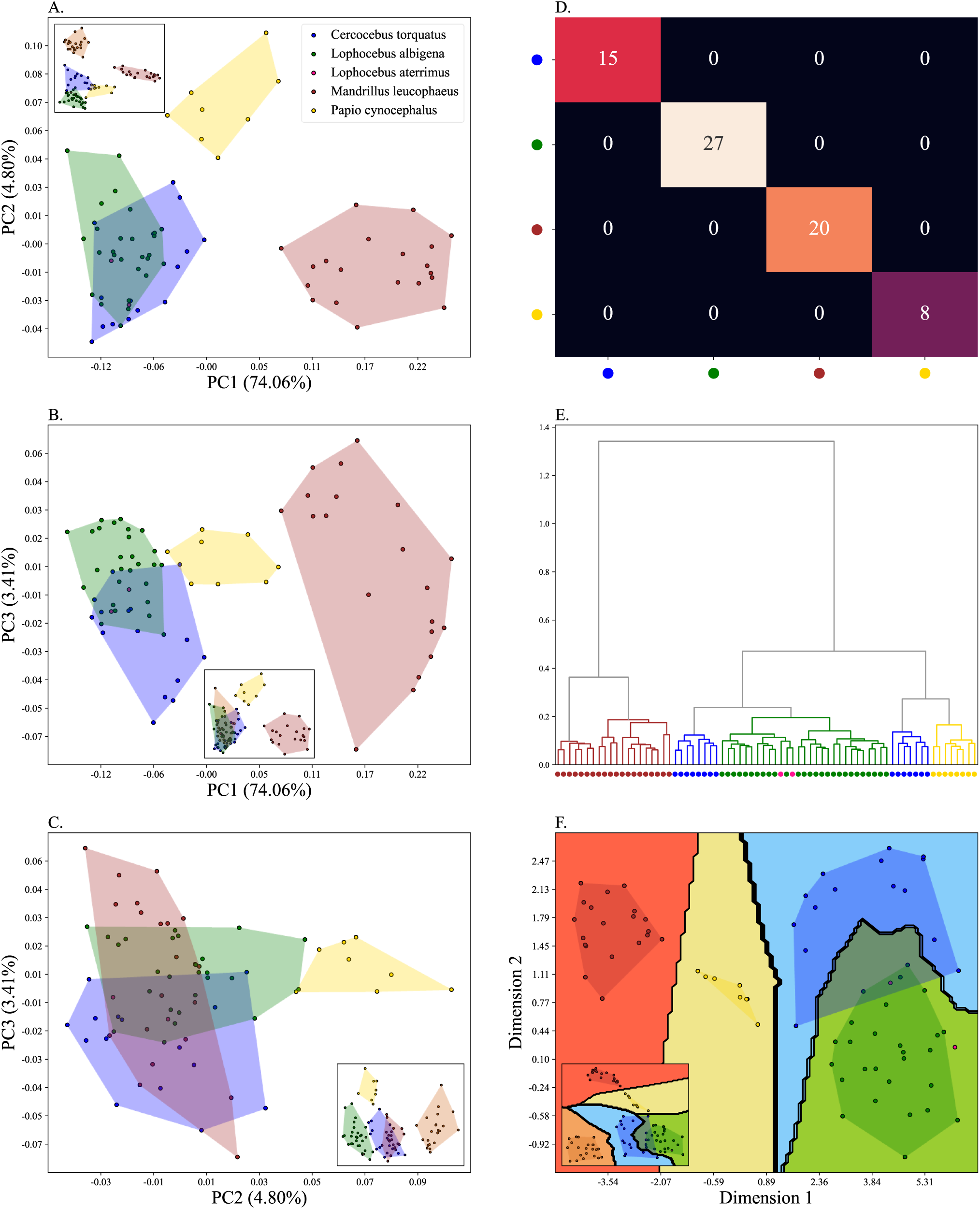
Plots depicting the papionin data after removing the Macaca mulatta (orange) taxon. A) PC 1-2, B) PC 1-3, C) PC 2-3, D) Confusion matrix of a 2NN classifier with 5-fold CV, E) Dendrogram of Procrustes distances of the samples using agglomerative clustering, F) Decision boundary of the 2NN classifier after t-SNE embedding with 5-fold CV. The insets in the subfigures are the corresponding benchmark PC scatterplots (Figure 4A-C).

Such results promote various forms of misclassifications and misinterpretations. For example, the single *Cercocebus torquatus (blue)* sample that falls inside the convex hull of the *Macaca mulatta (orange)* (Figure 4A) previously clustered with other samples from its own taxon (Figure 4A) and now clusters with *Macaca mulatta (orange)*. Misinterpretations of evolutionary processes, such as gene flow and phenotypic variations, can also be made regarding this sample. This *Cercocebus torquatus (blue)* sample could be claimed to belong to a taxon related to *Cercocebus torquatus (blue)* and *Macaca mulatta (orange)* experiencing gene flow. It could also be narrowed down to being their hybrid. Another notable example is the supposed relationships between *Cercocebus torquatus (blue)*, *Papio cynocephalus (yellow)*, and *Macaca mulatta (orange)* perceived from different plots. Figure 4A implies an old event of admixture between *Papio cynocephalus (yellow)* and *Macaca mulatta (orange),* which resulted in the *Cercocebus torquatus (blue)* with the appearance of breeding barriers separating it from both. Such a trend cannot be observed in the benchmark; By contrast, an opposite trend is obvious in Figure 4A, which implies a more recent event of speciation between *Cercocebus torquatus (blue)* and *Papio cynocephalus (yellow),* both being distanced from *Macaca mulatta (orange)*.

The changes in PC plots are even more profound in the case of *Macaca mulatta (orange)* removal. Here, *Cercocebus torquatus (blue)* and *Lophocebus albigena (green)* clusters almost overlap entirely across all the PC plots (Figures 6A -6C), whereas their overlap was minimal in the benchmark plot (Figure 4A). While the benchmark implies that the evolutionary distance between them is considerably high due to the accumulation of genetic differences to the point of speciation, Figure 6A is indicative of much higher genetic relatedness. The presence of only five *Lophocebus albigena (green)* samples not overlapping *Cercocebus torquatus (blue)* (Figure 6A) may be perceived as uniqueness within the genera; at the same time, the possibility of a speciation event cannot be ruled out. Considering the distribution of *Cercocebus torquatus (blue)* and *Lophocebus albigena (green)* over PC3, Figures 5B and 5C may imply an opposite trend, that the genetic divergence between the two is high, which can be indicative of penultimate states of speciation, especially if based on the location of the excavations, geographic isolation, and breeding barriers could be proposed. We note that the EV of these plots (Figures 4A and 5A) is higher than that of the benchmark (Figure 3A) since there are fewer specimens, implying a lower dimensional shape space, supposedly lending them more credibility by the widely assumed interpretation.

The overlap and separateness of taxa over PCs are frequently associated with morphological traits and trends. Methods useful for the study of morphometrics are expected to maintain an accurate representation of the relationships of the studied taxa and remain, at least largely, unaffected by the addition or removal of a taxon. In the case of PCA, we observed the exact opposite; while in the benchmark, *Macaca mulatta (orange)* and *Mandrillus leucophaeus (red)* are completely separated by PC2 (Figures 4A and 4C) after the removal of *Lophocebus albigena (green)*, *Macaca mulatta (orange)* overlaps considerably with *Mandrillus leucophaeus (red)* (Figures 5A and 5C). Thereby, it could either be claimed that PC2 *separates well* between the *Macaca mulatta (orange)* and *Mandrillus leucophaeus (red)* (Figures 4A to C) or that it is *taxonomically noninformative* about them (Figures 5B and 5C). These results are unsurprising because adding or removing specimens to PCA may result in a rotation of the PCs and reflect completely different portions and aspects of shape variation with different interpretations of the new space. The extent of this bias, shown here for the first time, demonstrates that PCA is a hazardous and unpredictable tool for studying morphological trends.

Concerning the use of explained variance as a measure of accuracy or reliability, it is notable that Figures 4A and 5A each explain more than 75% of the variance in shape (very close to that of the benchmark). Yet, the distances between specimens/species and, consequently, interpretations based on them greatly diverge. Moreover, classifying the samples using PC plots would lead to the misclassification of many in the case of *Macaca mulatta (orange)* removal. For instance, in Figure 6A (explained variance = 78%), at least 20 samples would potentially be misclassified.

Of the eight classifiers applied to the data, three (NN, GP, and SVC) and four (NN, LR, GP, and SVC) reached perfect accuracy (100%) in classifying the samples in a five-fold stratified CV in the first and second test, respectively. Similar to the benchmark, NN, LR, GP, and SVC show superior performance to the tree-based methods in the two cases of taxon removal. The LDA of the first two PCs had a considerably lower performance than other classifiers (accuracy = 80% in the second case), while the LDA of the first 10 PCs had perfect accuracy (100%) in both tests. The t-SNE plot creates well-separated clusters (Figures 5F and 6F), and the simple 2NN classifier has no difficulty differentiating groups (Figures 5E and 5F and 6E and 6F).

### The case of missing samples

To examine the effect of missing specimens on PCA and the alternative methods, we removed 11 and 19 samples from one (first test) and two (second test) taxa, respectively. As expected, we found that the shape and the area of the convex hull change with respect to the benchmark, specifically, that of the taxon(a) from which the samples were removed. This reduction is evident in all the PC scatterplots (Figures 7A-C and 8A-C). More concerningly, we observed changes in the clusters that may lead to misclassification of the unaltered taxa. For example, we note one of the two *Lophocebus aterrimus (pink)* samples (marked with a red arrow) in Figures 7A and 7C and 8A and 8C, which clustered with *Lophocebus albigena (green)* close to the *Cercocebus torquatus (blue)* in the benchmark (Figure 4A) changed its position. In Figure 7A, it is positioned close to the *Cercocebus torquatus (blue),* while in Figure 8A, it is outside the *Lophocebus albigena (green)* convex hull, far from other taxa. In both Figures 7C and 8C, it falls in the space between the *Cercocebus torquatus (blue)* and *Lophocebus albigena (green)*. Other inconsistent outcomes based on the PC plots were also evident; for example, the *Cercocebus torquatus (blue)* and *Mandrillus leucophaeus (red)* proximity varies in PC2 (Figures 4C, 7C, and 8C). In the benchmark (Figure 4C), the two groups largely overlap across PC2. By contrast, in Figure 7C, these taxa are largely distinct. Finally, Figure 8C implies that the taxa are completely separated. The interpretation of these relationships depends on the specific dataset. In the benchmark (Figure 4C), the two groups largely overlap in PC2 to the extent that PC2 may be considered *noninformative*; this overlap could also be inferred as a strong similarity in the genetic content implying a short evolutionary distance, i.e., the difference between the two taxa suggests *taxonomical identity*. By contrast, in Figure 7C, these taxa are largely distinct, which can be attributed to multiple possible factors, including the potential occurrence of divergence subsequent to the accumulation of sufficient genetic differences. This may also suggest the presence of breeding boundaries. Finally, Figure 8C implies that the taxa are completely separated. In this case, PC2 appears to separate all taxa, and speciation could be concluded here. Interestingly, this figure has the lowest explained variance (12.5%). Common to all these interpretations is that they are equally correct from a mathematical perspective, as they were all generated using the same method. However, they offer biologically conflicting interpretations, indicating the inappropriate use of PCA for morphometrics.

**Figure 7:**
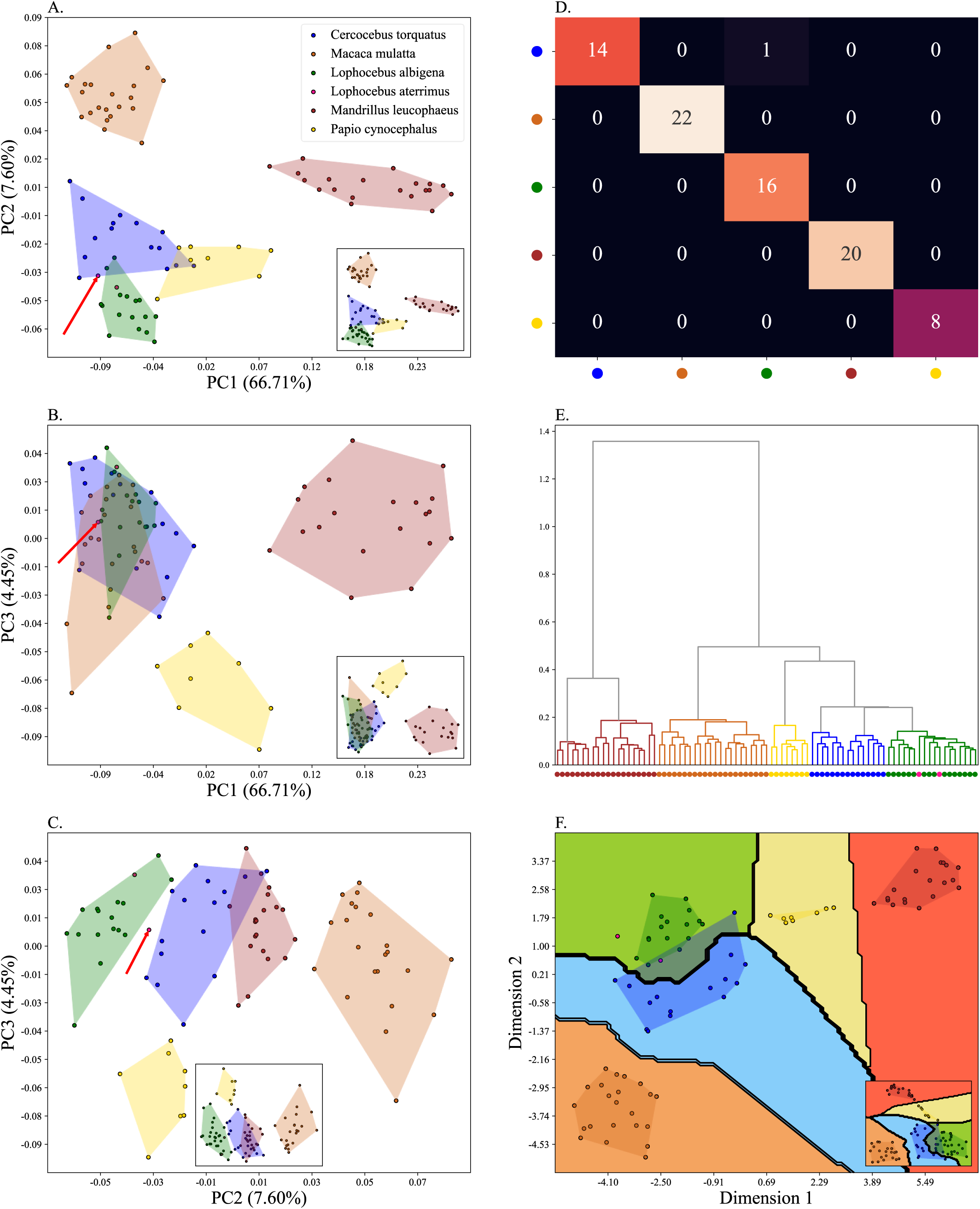
Plots depicting the papionin data after the first case of sample removal. 11 *Lophocebus albigena (green)* samples (index in benchmark dataset: 52, 56, 60, 63, 64, 66, 67, 73, 75, 76 and 77). A) PC 1-2, B) PC 1-3, C) PC 2-3, D) Confusion matrix of a 2NN classifier with 5-fold CV, E) Dendrogram of Procrustes distances of the samples using agglomerative clustering, F) Decision boundary with 5-fold CV. The insets in the subfigures are the corresponding benchmark PC scatterplots (Figure 4A-C). The red arrow in subfigures A, B, and C marks the *Lophocebus albigena (pink)* sample whose position in PC scatterplots is of interest.

**Figure 8:**
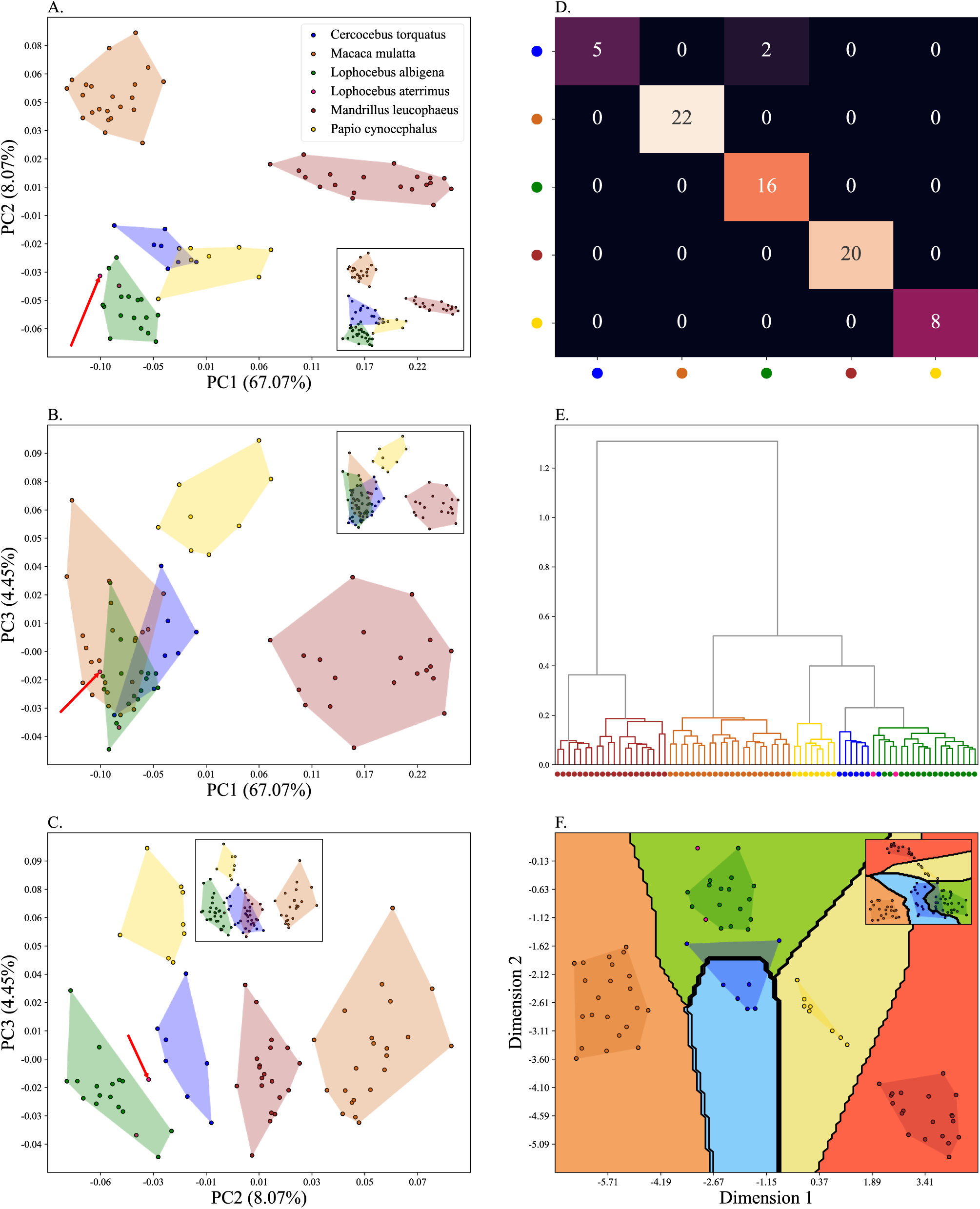
Plots depicting the papionin data after the second case of sample removal. 19 samples from both *Lophocebus albigena (green)* (index in benchmark dataset: 52, 56, 60, 63, 64, 66, 67, 73, 75, 76 and 77) and *Cercocebus torquatus (blue)* (index in benchmark dataset: 80, 81, 82, 83, 84, 88, 91 and 94) A) PC 1-2, B) PC 1-3, C) PC 2-3, D) Confusion matrix of a 2NN classifier with 5-fold CV, E) Dendrogram of Procrustes distances of the samples using agglomerative clustering, F) Decision boundary with 5-fold CV. The insets in the subfigures are the corresponding benchmark PC scatterplots (Figure 4A-C). The red arrow in subfigures A, B, and C marks the *Lophocebus albigena (pink)* sample whose position in PC scatterplots is of interest.

Investigating the classifiers proved to be considerably robust against these types of omissions. For the first test, all the classifiers apart from XGBoost reached accuracies equal to 96% and higher. For the second test, which is more challenging, three classifiers (NN, GP, and SVC) have accuracies greater than 94% and the remaining range between 85% and 91% in the aforementioned CV scheme (Table 2). Compared to other classifiers, the LDA of the first two PCs exhibited poorer performance, whereas the LDA of the first 10 PCs achieved perfect accuracy (100%) in both tests. It can be noted that the 2NN classifier had difficulties in classifying two *Cercocebus torquatus (blue)* samples because, due to the removal of samples from their own taxon, their nearest neighbours were now *Lophocebus albigena (green)* (Figure 8F).

**Table 2:** The LOF outlier scores. *Cercocebus torquatus (blue)* was treated as an outlier in three cases; A) each sample separately, B) 6 samples together, and C) the whole group together.

| Sample | A) LOF score<br>(a) | B) LOF score<br>(b) | C) LOF score<br>(c) |
| --- | --- | --- | --- |
| 80 | -1.22 | - | -0.98 |
| 81 | -1.21 | - | -1.02 |
| 82 | -1.16 | - | -1.01 |
| 83 | -1.34 | - | -0.99 |
| 84 | -1.36 | -1.07 | -1.03 |
| 85 | -1.27 | -1.03 | -1.03 |
| 86 | -1.21 | -1.01 | -1.02 |
| 87 | -1.24 | -1.12 | -1.06 |
| 88 | -1.13 | - | -1.04 |
| 89 | -1.22 | - | -1.06 |
| 90 | -1.32 | -1.00 | -1.01 |
| 91 | -1.08 | - | -1.03 |
| 92 | -1.25 | -1.07 | -1.08 |
| 93 | -1.37 | - | -1.24 |
| 94 | -1.19 | - | -1.13 |

### The case of missing landmarks

Newfound samples often comprise incomplete osteological remains or fossils (18, 22) and only present partial morphological information. Choosing landmarks is an important and controversial subject (2, 23, 24). However, the ability of a landmark set to predict the shape of the incomplete part is questionable. To evaluate this effect in downstream analyses, we studied two cases of landmark removal, each time removing one set (comprising all three cartesian coordinates of each landmark). First, we removed one set (Figure 9) with the symmetric pairs of the omitted landmarks remaining (Table S4). Next, we removed another set, of which more than half had no symmetric pairs (Figure 10).

**Figure 9:**
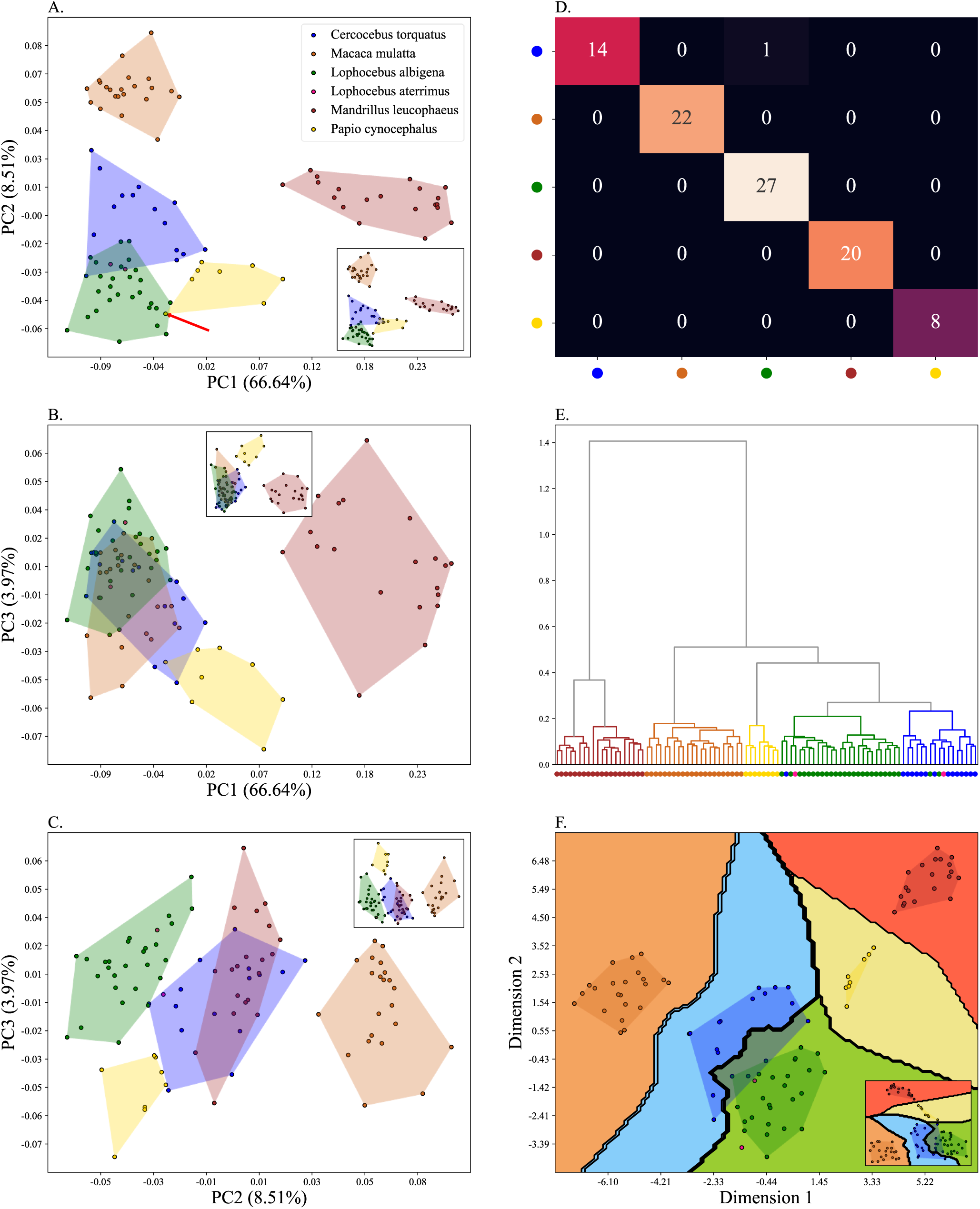
Plots depicting the papionin data after the first case of landmark removal. Landmarks 23, 24, 25, 26 and 27 were removed. Scatterplots of the papionin data after the first case of landmark removal. A) PC 1-2, B) PC 1-3, C) PC 2-3, D) Confusion matrix of a 2NN classifier with 5-fold CV, E) Dendrogram of Procrustes distances of the samples using agglomerative clustering, F) Decision boundary with 5-fold CV. The insets in the subfigures are the corresponding benchmark PC scatterplots (Figure 4A-C).

**Figure 10:**
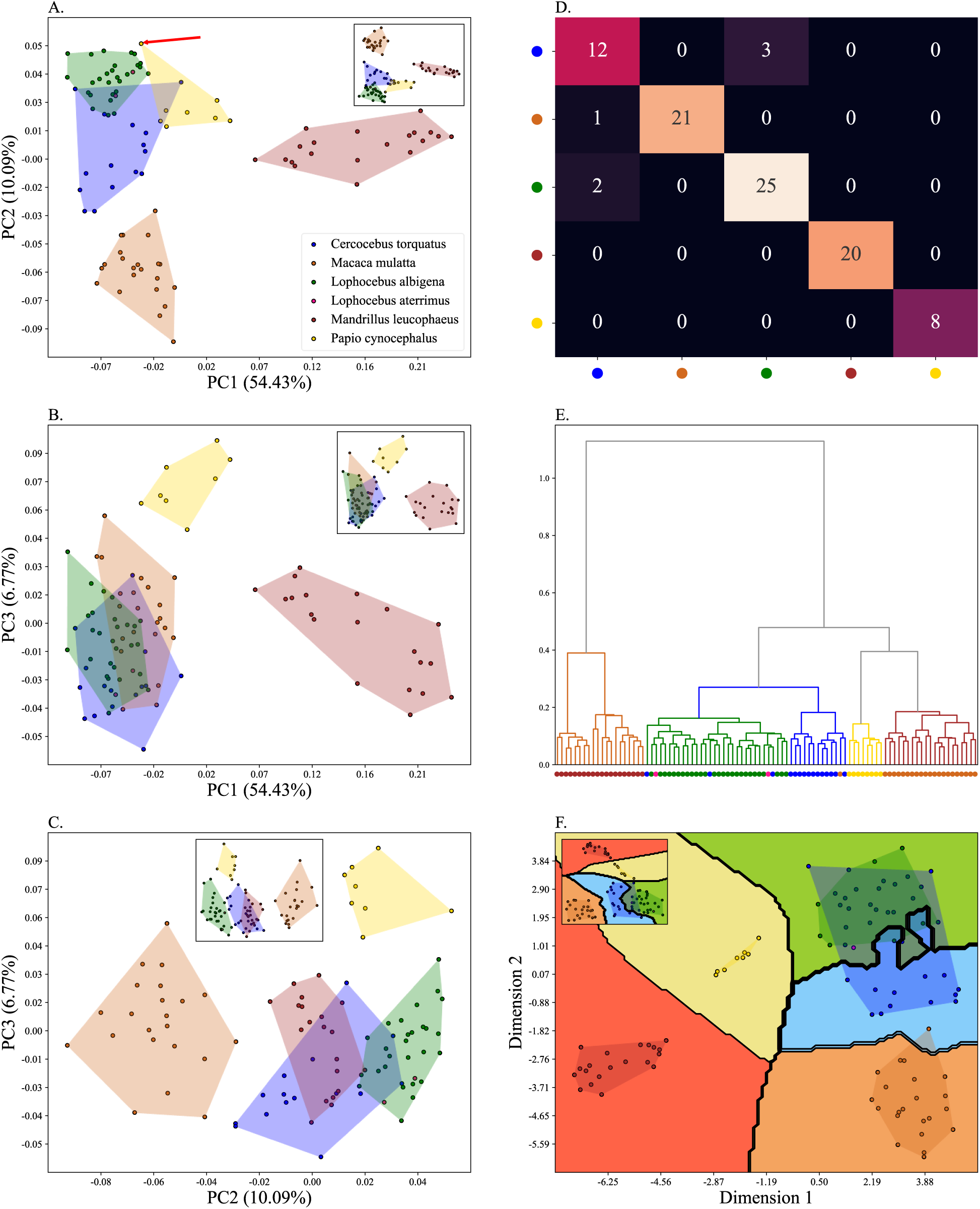
Plots depicting the papionin data after the second case of landmark removal. Landmarks 11, 12, 16, 17, 18, 30 (symmetrical pair of 12) and 31 (pair of 13) were removed. PC 1-2, B) PC 1-3, C) PC 2-3, D) Confusion matrix of a 2NN classifier with 5-fold CV, E) Dendrogram of Procrustes distances of the samples using agglomerative clustering, F) Decision boundary with 5-fold CV. The insets in the subfigures are the corresponding benchmark PC scatterplots (Figure 4A-C).

This type of alteration results in deviations from the benchmark in the shape of the convex hulls and the distances between the samples and taxa clusters across all the PCs (Figures 9A to 9C and 10A to 10C). For brevity, we only point out the most distinct outcomes. First, we investigate the perceived evolutionary relationships between *Papio cynocephalus (yellow)*, *Lophocebus albigena (green)*, and *Cercocebus torquatus (blue)* (Figures 4, 9, and 10). In the benchmark (Figure 4A) *Cercocebus torquatus (blue)* slightly overlaps with *Lophocebus albigena (green).* This overlap increases considerably in both Figures 9A and 10A. Moreover, by contrast to the benchmark where the *Papio cynocephalus (yellow)* and *Lophocebus albigena (green)* are completely separated, in Figure 9A, one single *Papio cynocephalus (yellow, marked with a red arrow)* sample (index in the benchmark: 6) now clusters with *Lophocebus albigena (green)*, raising challenges in identifying its origins and relatedness to other taxa. The position of the same *Papio cynocephalus (yellow, marked with a red arrow)* sample in Figure 10A is also notable. After removing the second set of landmarks, it appears as an outlier. Hence, a range of gene flow hypotheses and other possible causes of shape difference between this sample and both the *Papio cynocephalus (yellow)* and *Lophocebus albigena (green)* can be proposed. The position of the same *Papio cynocephalus (yellow)* sample in Figure 9A is also notable. After removing the second set of landmarks, it appears as an outlier. Thereby, while the complete skull would cluster with its own taxon, with only a small portion of its zygomatico-maxillary region missing, it could either be misclassified as *Lophocebus albigena (green)* (Figure 9A) or considered an outlier, possibly belonging to a new taxon (Figure 9A).

Another notable inconsistency is the relationship between *Cercocebus torquatus (blue)*, *Lophocebus albigena (green)*, and *Mandrillus leucophaeus (red)* over PC2, best seen in PC2- PC3 scatterplots (Figures 4C, 9C, and 10C). In the benchmark and the first case of landmark removal, they are *well separated* by that PC (Figures 4C and 9C, respectively); however, in Figures 10C *Lophocebus albigena (green)* and *Mandrillus leucophaeus (red)* are tangent over PC2. Accordingly, Figures 4C and 9C suggest considerably more evolutionary distance compared to Figures 10C. Different interpretations can be proposed for these plots. Additionally, in Figures 4C and 9C, *Cercocebus torquatus (blue)* appears only to exchange genetic material with *Mandrillus leucophaeus (red)*. However, in Figure 10C, the exchange was between *Lophocebus albigena (green)* and *Mandrillus leucophaeus (red)*. This, in turn, can be interpreted as a loss of biodiversity due to hybridisation (62, 63, 64, 65). Similar scenarios can be observed when feral and domesticated species mate (66). This may lead to different interpretations concerning the population dynamics depending on factors such as the frequency of such events, the population size, and the selection pressures. Consequently, this could even lead to homogenization (67).

Considering the supervised classifiers, this type of alteration resulted in a negligible drop in accuracy (Tables 1 and 2). However, their performances were acceptable in both cases. Three (NN, GP, SVC) and two (NN, GP) had accuracies above 97% and equal to 95% in the first and second cases, respectively, higher than the LDA of the first two PCs. The LDA of the first 10 PCs performs very well in both tests (accuracy = 100% and 96% in the first and second tests, respectively). Due to distortions in the second case, some of the *Cercocebus torquatus (blue)* and *Lophocebus albigena (green)* samples have moved close to the other taxon and were misclassified by the 2NN classifier (Figures 10F). In the first case, the accuracies of RF and ET were above 90%, while in the second case, they dropped below 90%. With an accuracy below 85%, XGBoost had the lowest performance. The results indicate the robustness of the classifiers to partial missingness in landmarks.

### The case of combined alterations

Thus far, for simplicity, we have introduced and studied the three types of alterations separately as if they represent separate challenges associated with ancient fossils; however, in reality (18, 22, 68), when a fossil is found, it might only be a fragment (fragments) of a bone (landmark removal) incomparable with some fossils (sample removal). It is also common to compare samples of interest with much larger cohorts of other taxa. For example, Harvati et al. (27) compared 12 Neanderthal specimens from Europe and the Near East; the early Neanderthal specimen from Reilingen, the Middle Pleistocene African specimen Kabwe, two early anatomically modern humans (AMH) from the Near East, and four Late Palaeolithic European AMHs and 270 modern humans from nine populations using fifteen temporal landmarks. In the case of *Homo NR*, Hershkovitz et al. (22) analysed four landmarks and 77 surface semi-landmarks in NR-1, 18 landmarks and 77 curved semi-landmarks in NR-2, and a set of six landmarks, 18 semi-landmarks (enamel–dentine junction (EDJ)) and two landmarks and 18 semi-landmarks (cemento-enamel junction (CEJ)) in the lower left second molar in a combined analysis of the junctions. In each analysis, the sample was compared with a different set of humans. In both studies (22, 27), the number of samples representing each group was utterly dissimilar; some groups were not presented, and some were under/over-presented. In the case of *NR* fossils (22), the use of only four landmarks (specifically on the cranium) was another limitation (22). Unfortunately, despite numerous appeals to the authors and journals, neither group released their complete data publicly, thus hindering reproducibility.

Next, we studied how these combined limitations affect morphometric analyses using two test cases. In the first case, one taxon (*Mandrillus leucophaeus, red*) and a greater portion of the collective samples were randomly removed (30/94 samples remained), and only a portion of the cranium (landmarks #1, 2, 3, 4, 6, and 14) was investigated. In the second case, all the taxa were present, but only a few samples (37/94) and a small portion of the cranium (landmarks #1, 2, 3, 4, and 14) were kept. Figures 11 and 12 show the results of the PCA and the 2NN classifier for the two cases, respectively. At odds with the benchmark (Figure 4A), in the PC scatterplots, many groups overlap, and the samples are poorly separated. Unsurprisingly, the performance of the ML classifiers is considerably low for these tests (Tables 1 and 2), with balanced accuracies slightly above 50% and 40% for the first and the second cases, respectively. In such cases, the interpretability of the data is questionable as a fundamental lack of information in the dataset will bias all methods. In other words, the typical landmark datasets in palaeoanthropology analysed using the established pipeline (Figure 2) yield incorrect results that can only be partially improved with alternative methods.

**Figure 11:**
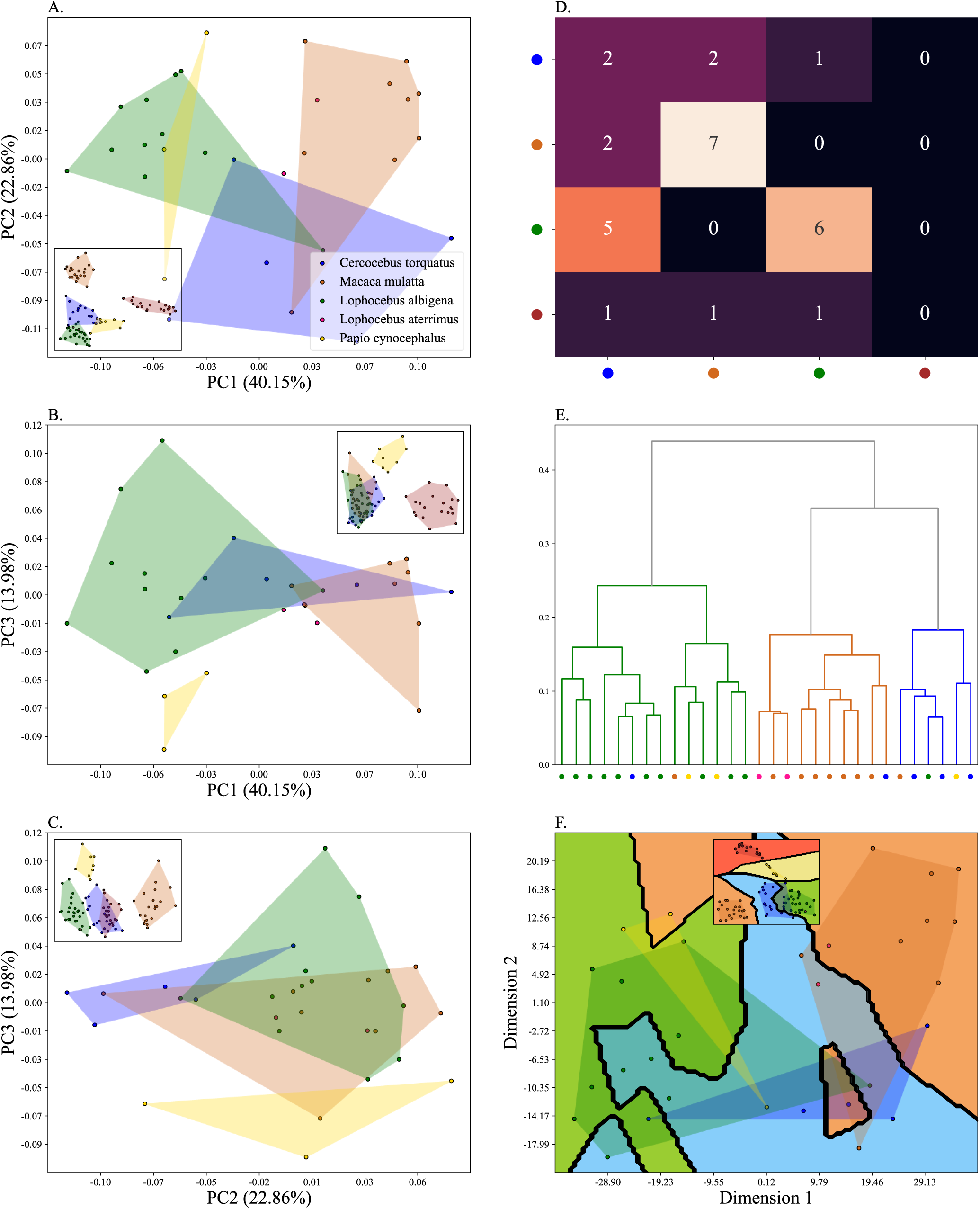
Plots depicting the papionin data in the first extreme case. A) PC 1-2, B) PC 1-3, C) PC 2-3, D) Confusion matrix of a 2NN classifier with 5-fold CV, E) Dendrogram of Procrustes distances of the samples using agglomerative clustering, F) Decision boundary with 5-fold CV. The insets in the subfigures are the corresponding benchmark PC scatterplots (Figure 4A-C). the corresponding benchmark PC scatterplots (Figure 4A-C).

**Figure 12:**
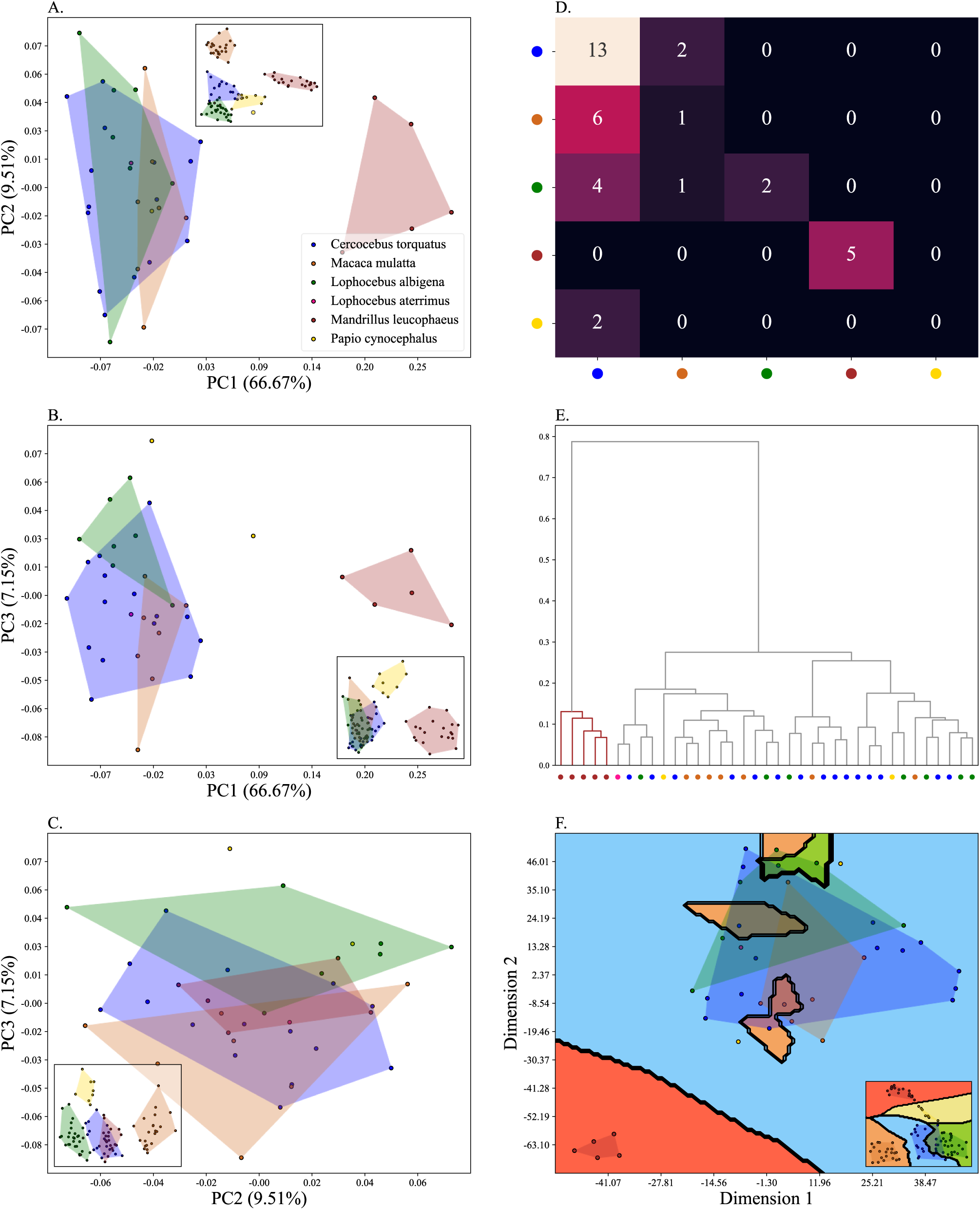
Plots depicting the papionin data in the second extreme case. A) PC 1-2, B) PC 1- 3, C) PC 2-3, D) Confusion matrix of a 2NN classifier with 5-fold CV, E) Dendrogram of Procrustes distances of the samples using agglomerative clustering, F) Decision boundary with 5-fold CV. The insets in the subfigures are the corresponding benchmark PC scatterplots (Figure 4A-C).

### The case of outlier detection in a specimen of unknown taxonomy

Newfound samples may represent individuals (18, 22, 68) of unknown taxa (18, 42, 68). Therefore, identifying them requires a different approach than classification because classifiers can only classify samples to already known classes and, thus, are not capable of detecting outliers (taxa). To address this issue, we investigated the ability of three methods to detect outliers by measuring the outlier score for outlier samples using the Local Outlier Factor (LOF) (69), isolation forest (70), and one class SVM (71) since the isolation forest and one class SVM methods yielded poor results compared to LOF (Figures S8 and S9) we did not investigate them further.

When exploring the origins of unidentified samples through morphometrics, it is crucial to exercise extreme caution. As early as 1978, Oxnard (19) emphasised the necessity for meticulous attention, particularly when examining a single specimen, more so in the case of fossils, where researchers may unconsciously seek to validate their intuitive speculations using analytical methods. To address this and investigate the performance of outlier detection methods, we carried out three trials by considering *Cercocebus torquatus (blue)* samples as novelties (outliers) and calculated their outlier score in all tests. First, to address the case of finding a single (22) specimen, we added each *Cercocebus torquatus (blue)* sample to the cohort separately. Second, in the case of several (68) individuals belonging to the same taxon, we performed two trials; we added together six *Cercocebus torquatus (blue)* in each other’s proximity. In the last trial, we added 15 *Cercocebus torquatus (blue)* samples to the dataset. Figures 13-15 depict the results. Eq. 1 yields the circle’s radius around the *i_th* sample in the scatterplots. The LOF scores for each sample in all three tests are presented in Table 2. When treated separately, samples receive considerably high scores, identifying them as outliers. As expected, adding samples together reduces their outlier scores (smaller magnitude, greater negative score) because LOF evaluates the samples based on the concept of local density, and samples from one group are expected to be in a neighbourhood in the shape space manifold (considering that the data is informative). If the LOF scores are reduced after being treated together, then it could mean that they may belong to an as-of-yet-unknown group. These results confirm the ability of LOF to identify novelties in landmark data shape space.

**Figure 13:**
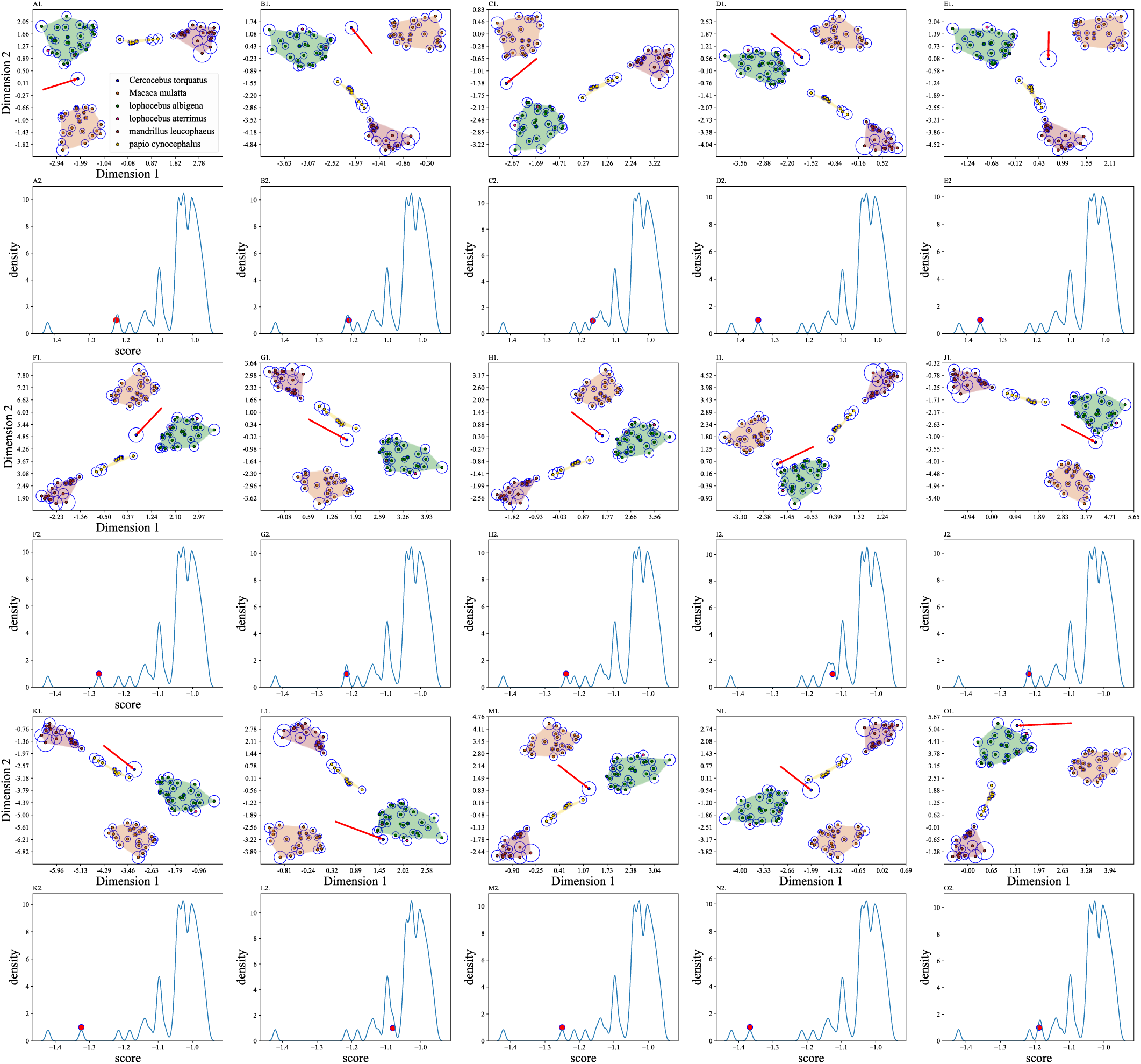
Detecting outliers using local outlier score and KDE plots. Each *Cercocebus torquatus (blue)* sample is treated as an outlier (red arrow, A1-O1 t-SNE plots) and the KDE of the scores are plotted for each case (A2-O2). The radius of the circle around each sample is proportional to the LOF score. The red dot in the KDE plots shows the LOF score of the outlier in each case.

**Figure 14:**
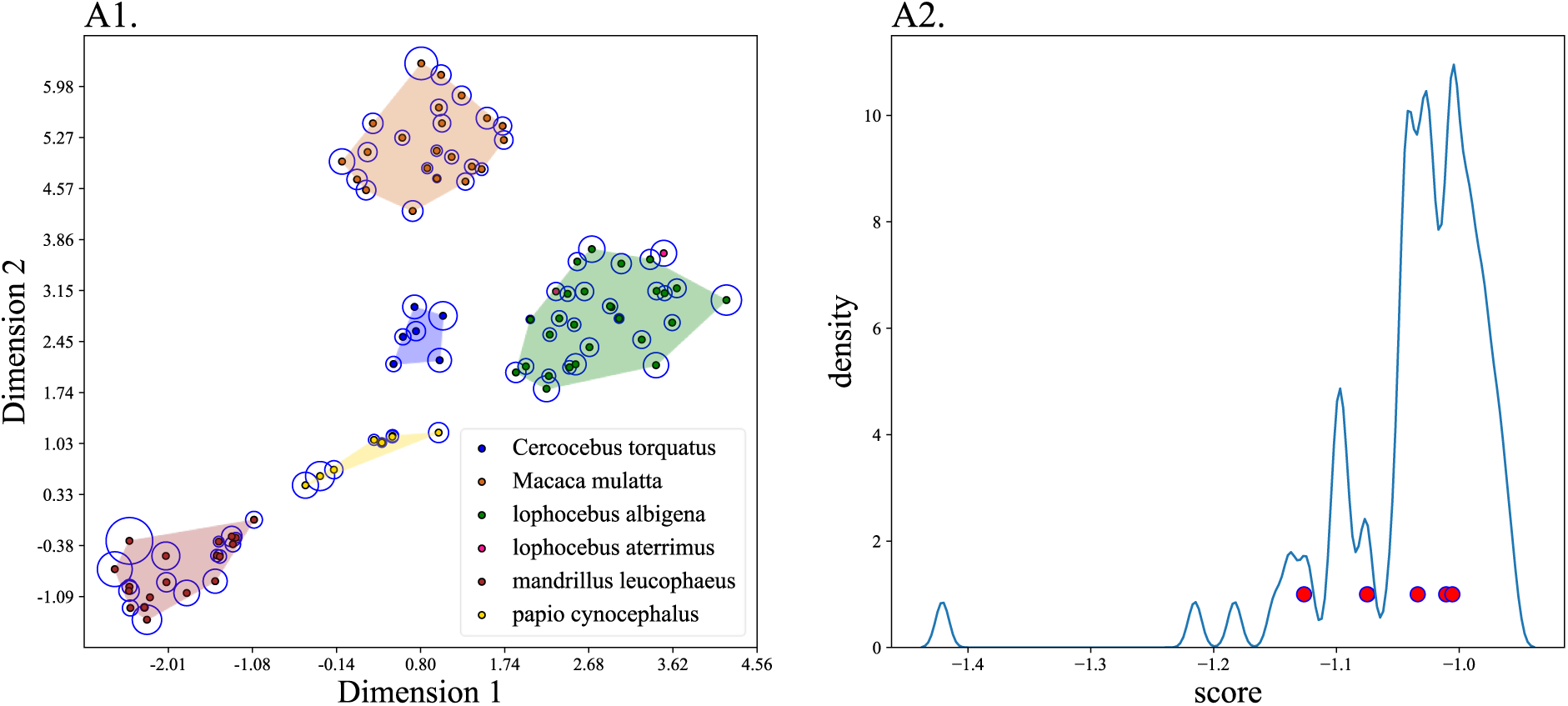
Detecting outliers using local outlier score and KDE plots. Six *Cercocebus torquatus (blue)* samples (indexes: 84,85,86,87,90 and 92) are treated as outliers (A1 t-SNE plots) and the KDE of the scores are plotted (A2). The radius of the circle around each sample is proportional to the LOF score. The red dots in the KDE plots show the LOF score of the outliers.

**Figure 15:**
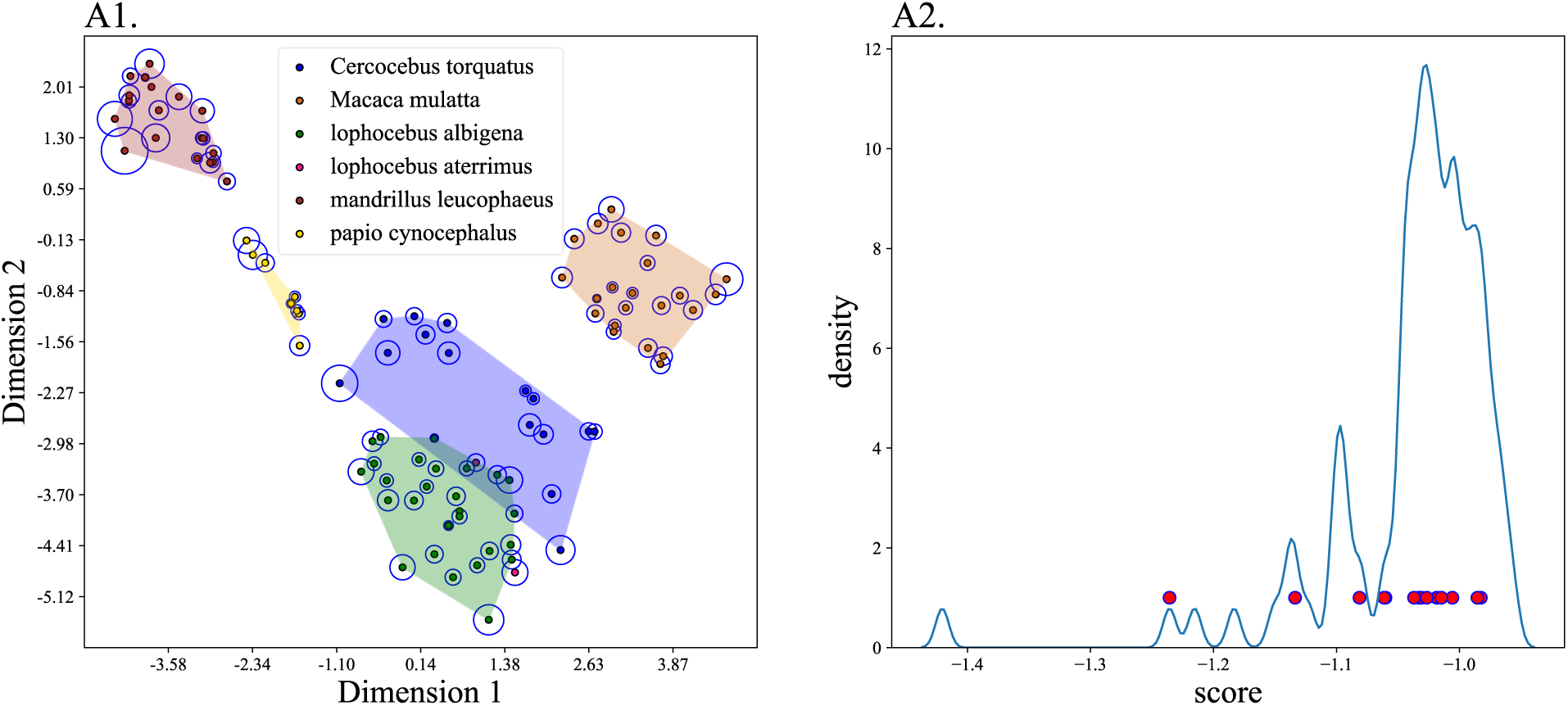
Detecting outliers using local outlier score and KDE plots. All *Cercocebus torquatus (blue)* samples are treated as outliers (A1 t-SNE plot) and the KDE of the scores are plotted (A2). The radius of the circle around each sample is proportional to the LOF score. The red dots in the KDE plots show the LOF score of the outliers.

**Figure 16:**
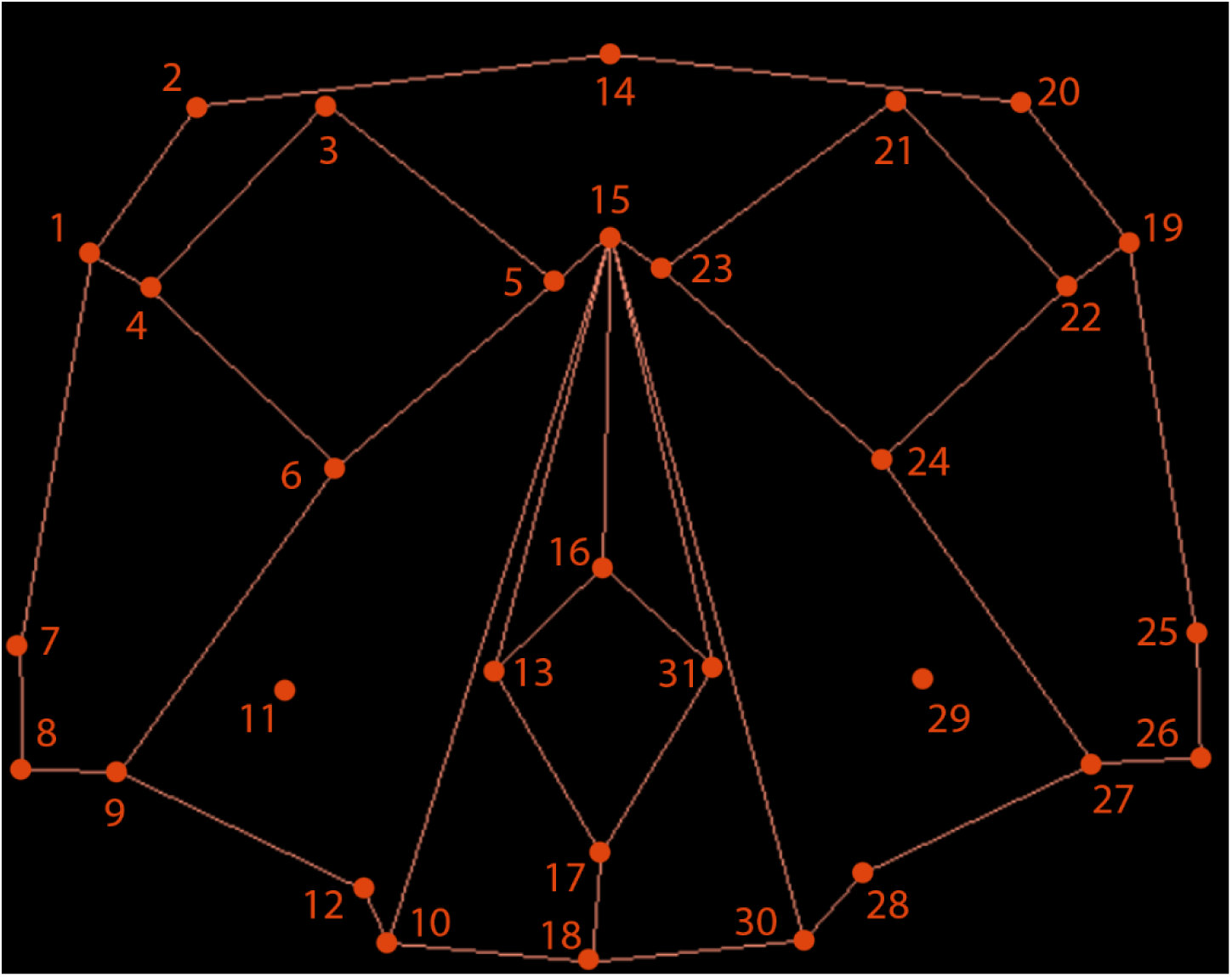
The 31 landmarks of Table S1. The wireframes connect the landmarks.

### Evaluation of the informativeness of landmarks with feature selection

Using informative input is vital to the success of ML-based classification methods. It is well established that the required number of data points for an accurate outcome is (generally) exponentially proportional to the number of data dimensions (72), a phenomenon that Bellman termed the “*curse of dimensionality*” (73). In other words, an analysis feasible in low-dimensional spaces can be challenging or meaningless in higher dimensions as data becomes too sparse (74). To overcome this problem, it is necessary to reduce the number of dimensions through either feature selection methods that select informative features (75) or feature extraction/transformation methods that create new features from the original variables (76). To assess the informative or noninformative features, we carried out several iterations of feature selection, although the low number of landmarks in our benchmark dataset did not handicap the classifiers (Tables 1 and 2). For that, we applied a genetic algorithm (77, 78, 79, 80) as a wrapper method around a logistic regression (81) classifier five times and calculated the accuracy of the classifier with the identified subsets in each case (Tables S2 and S3). Despite considerable differences in the identified landmark coordinate subsets, all subsets performed with similar accuracy, nearly equal to the original set, although half its size. In other words, half of the landmark coordinates are either redundant, noninformative, or both.

## Discussion

The quantitative analysis of morphological data has always captivated researchers (3, 4). Employing graphical and computational methods, as done today, is rooted in ancient geometric principles and developments since the European Renaissance (1, 2). However, it was not until the mid-1990s (6, 7, 8, 20) that the mathematical underpinnings of shape found solid ground, and complemented by cutting-edge graphics and statistical tools, propelled modern morphometrics into the realm of essential tools spanning diverse fields, including evolutionary biology, anthropology, palaeontology/palaeoanthropology, and even forensics (9, 14, 15, 16, 17). In biological science, morphometrics is used to study temporal shape variation inter- and intra-species and is instrumental in exploring questions – from the intricate changes in shapes during evolution to questions concerning growth and development. There are various approaches in morphometrics, but among them, geometric morphometrics has left an indelible mark on biology, especially in anatomy and physical anthropology.

Geometric morphometrics encompasses a diverse range of approaches; most share a standard workflow involving two core steps: Generalised Procrustes Analysis (GPA) followed by Principal Component Analysis (PCA) (Figure 1) (2, 16, 17, 25, 26). GPA aligns landmark coordinates by minimising the shape-independent variations across samples. PCA projects the data onto uncorrelated variables called principal components (PCs), which capture different proportions of variation within the data (13). The PCs are then plotted on colourful scatterplots and examined visually. Generally, biologically related shapes are expected to exhibit closer clustering, while unrelated shapes will be more distanced and dispersed. However, there are no rules on which plots to use. Researchers typically select favourable PCs, disregarding the remaining ones (22, 27, 28, 29), and claim that various PCs, i.e., statistical constructs agnostic to the underlying data (37), represent demographic and evolutionary processes (18, 27, 32). Interpretations may also encompass origins, relatedness, gene flow, speciation, phenetic evolutionary trends, and phenotypic/genotypic variation within samples and taxa (16, 17, 18, 22, 26, 27, 28, 29, 32, 33, 34). Additional assumptions include that variations or covariations along specific PCs are often linked to specific morphological traits (13, 22, 36) and that PCs with *sufficiently higher explained variances (EVs)* represent certain phenotypic (2, 22, 27, 32) or even sexual (27, 32) morphological traits.

Given the widespread popularity of PCA in geometric morphometrics (18,400-35,200 studies), researchers and practitioners may naturally anticipate that it will yield accurate and valid results. The growing body of literature criticising the accuracy of various PCA applications (47, 48, 51, 82, 83, 84) raises concerns about its accuracy in geometric morphometrics. Surprisingly, despite its far and wide applications, PCA has never been shown to produce *correct* shape inferences. For that, we undertook the first thorough investigation of the possible shortcomings of PCA in geometric morphometrics, where we assessed its reliability, robustness, and reproducibility in shape analysis compared to eight alternative ML methods. Specifically, we evaluated PCA performance for four use cases using a benchmark dataset, to which we introduced eight alterations simulating real-life scenarios. In the fifth use case, we investigated the accuracy of ML methods to perform outlier detection in the case of novelties, i.e., the identification of new taxa.

A key strength of our study is using benchmark data as the “ground truth” to assess the accuracy of PCA using t-SNE, dendrograms and confusion matrices as alternative visualisation aids. We found that the five papionin taxa cluster separately in the dendrogram of the shape space and its t-SNE plot (Figures 4E and 4F). This allowed us to evaluate the performances of PCA in light of alterations, representing real-life scenarios, which reflect the restraints faced in morphometric analysis of ancient samples (e.g., fossils). PCA scatterplots of the altered datasets changed the taxa’s positioning, the clusters’ shape, and the distances within and between clusters with respect to the benchmark. For example, samples from one cluster would overlap with others; clusters tended to appear more or less heterogeneous, and separate clusters would overlap. We showed that the PC scatterplots produce inconsistent results for the same dataset based on the choice of the PCs. We demonstrated how analysing different PCs gives different outcomes, which can lead to different conclusions (Figure 2). Using the *Homo Nesher Ramla* (*NR2*) as a test case (22), we demonstrated that the experimenter could influence the outcome by selecting preferable results out of all the conflicting options provided by PCA (Figure 2). Alternatively, researchers may see patterns emerging when analysing different PCs and uphold the most favourable ones. For example, Harvati (27), who analysed the *Skhul 5* (85), a 40,000-year-old human skull from *Mount Carmel* (Israel), proposed diverging hypotheses based on favourable PC outcomes (based on PC8 separating it from Neanderthals and modern humans and associating it with the Late Palaeolithic specimen and based on PC12 associating it with modern humans). Such an approach would lead to contradictory evolutionary misinterpretations, promoting misidentification and misclassification. Our results (Figures 5-12) show that compared to the benchmark (Figure 4), PCA produces results that are statistical artefacts of the input data, and although they can be interpreted in evolutionary terms such as gene flow, admixture, or speciation – they would nonetheless be artefacts and biologically meaningless. Overall, we found that PCA results are not reliable, robust, or reproducible as some members of the field assume (Table 1).

The popularity of PCA stands in sharp contrast with its multiple limitations, some of which were also reported in other fields (e.g., 51, 86). To name a few, PCA lacks any measurable power, a test of significance, or a null model, making it a problematic tool, at the very least. PCA outcomes are open to manipulations and interpretations, and the ability to evaluate various dimensions allows for cherry-picking of the results where all the answers are equally acceptable. Without external validation methods, such as those used by Marom and Rak (87) in their criticism of Hershkovitz et al.’s (22) work based on unique morphological characteristics of *H. neanderthalensis* mandible (e.g., the medial pterygoid tubercle), PCA-based arguments are vacuously true based on the antecedent of their correctness. It may be argued that PC results with “*sufficiently high explained variances”* should be favoured. No threshold for such a term exists, and we showed instances where higher explained variances in plots were less accurate than plots with lower explained variances (Figure S1). Overall, we found no justification for the use of PCA in morphometrics.

Beyond evaluating PCA’s accuracy, we comparatively investigated the accuracy of eight ML methods using the same use cases and benchmark in classifying superimposed landmarks, including LDA with the first two and 10 PCs. The ML classifiers showed superior performances across all the tests except the most extreme case of the combined alterations, where all the analyses produced inaccurate results (∼50% accuracy). These methods were more accurate in the first three use cases (80-100% accuracy) and less susceptible to experimenter intervention. It can be observed that the classifiers are also more stable with a decrease in the number of samples. Moreover, the low-dimensional embedding could form distinct clusters in the first three cases (Figures 5 to 10). Nonetheless, we note that ML methods also have limitations and are not immutable to datasets with high missingness rates, as demonstrated in the case of combined data alterations. Linear Discriminant Analysis (LDA) of the first two PCs exhibited relatively lower performance compared to the seven ML classifiers. Conversely, the LDA utilising the first 10 PCs demonstrated satisfactory performance. It is worth mentioning that while LDA is extensively utilised as the primary technique for classifying landmark data in the field, there is no established guideline regarding the optimal number of PCs to be incorporated in the Discriminant Analysis (DA). Moreover, a drawback of LDA applications is that they handle many variables (88). In practice, this selection is often made by the experimenter to obtain the preferable performances in a cross-validation (CV) (89) scheme and may include the first 30 (90) or 50 (91) PCs.

Of all the classifiers tested here, the simpler classifier (NN) was more efficient than the more advanced ML methods. Unsurprisingly, the tree-based classifiers had relatively low accuracies since they have a higher variance error component as they are more sensitive to data distribution. Tree-based algorithms partition the feature space based on the available data. When the sample size is small, the algorithm becomes more sensitive to the specific distribution of the training data; thereby, small variations or outliers in the data can substantially impact the resulting tree structure, leading to higher variance in predictions (92). The advantage of the NN classifiers as a substitute for PC scatterplots and to promote geometric morphometrics cannot be overstated.

The identification of novel taxa is always exciting – as long as it can be proven as such. Unsurprisingly, PCA has played a pivotal role in facilitating such “breakthroughs” (21, 54, 103, 117). To note one example, it has been assumed that craniofacial morphology can effectively serve as a population genetic proxy for investigating population structure based on its broader spatial and temporal coverage (100, 101, 116). It has further been claimed that this evidence is stronger than paleogenetic evidence based on ancient DNA derived from certain populations, such as inhabitants of the pre-Contact New World. Therefore, it was employed for multiple dispersal events and ancestral origins inferences (100, 101, 116). However, it has been proven conclusively that this does not hold true generally in disputes surrounding the case of the “Kennewick Man” (96). Moreover, it is important to note that palaeogenetics is based on at least thousands of biological markers with internal consistency, which can be effectively compared (93) as opposed to limitations faced in morphometric comparisons of ancient cranial remains.

Regrettably, challenges within the field of physical anthropology extend beyond the appropriate and justifiable use of statistical methods. It is often observed that datasets are not readily accessible, as they are privately held and not easily shareable. In an era that values open science and data-sharing (94), this system is maintained through collaborations of four main stakeholders, including authors, journals, funders, museums, and the media, to safeguard the survivability of physical anthropology and, by extension, their positions and interests. For example, Hershkovitz and May admitted that the co-authors of the Hershkovitz et al. paper (22) had signed a contract to keep the data private (personal communication), which nullifies the data sharing policy of Science. Science’s policy states that “the *Science* journals generally require all data underlying the results in published papers to be publicly and immediately available” (https://www.science.org/content/page/science-journals-editorial-policies). After falsely claiming that the data-sharing policy was not in place when the said paper was published, the Science editor upheld this violation of their own data-sharing policy. Once their unconstructive behaviour was pointed out, they ignored our further emails (personal communication). We next approached the funders who sponsored the study. Except for one funder, who claimed they did not fund the study despite being listed in the paper, our attempts to communicate with funders regarding this matter yielded limited responses. The funders either ignored our inquiries or indicated no concerns with the practice. The role of museums and the agreements they require scientists to sign when seeking access to morphometric information of samples should also be considered. On one end of the scale are museums willing to share data subjected to stipulated agreements, whereas on the other end are museums that adopt a more restrictive stance, particularly when dealing with valuable artefacts, and closely monitor who gains access to the data and how the information is utilised. Most museums lie somewhere between these two ends. Finally, the media that publish uncritical, exaggerated reports encourages researchers to use poor methods to satisfy this publicised promotion. For example, with respect to the *Nesher Ramla* fossils, The Times’s headline was: “Meet Nesher Ramla Homo: New form of human found,”(95) The Guardian’s headline was: “Fossilised bones found in Israel could belong to mystery extinct humans,”(96) and Science Alert’s headline was “A Previously Unknown Type of Ancient Human Has Been Discovered in The Levant.”(97) Remarkably, only a few sources, including Haaretz (98) and The Times (99) bothered to update their readers about the substantial controversy ensuing from the findings of Marom and Rak that the *Homo Nesher Ramla* is a Neanderthal (87), but this is not the general case. In the more recent case of HLD 6 (18), CNN’s headline was ‘300,000-year-old skull found in China unlike any early human seen before’ (100), and The Daily Mail’s headline was ‘Have scientists discovered a new species of HUMAN? Ancient skull belonging to a child with no chin who lived 300,000 years ago suggests our family tree ’needs another branch’’ (101). This was later echoed in a Nature News article titled “A new human species? Mystery surrounds 300,000-year-old fossil” (102).

Our findings and the widespread scientific misconduct we encountered in the field provoke severe concerns regarding a major body of morphometrics analyses. We observed that data are kept intentionally covet to avoid the falsifiability of evolutionary theories. When empirical tests cannot contradict a statement of empirical science, it is no longer a statement that uncovers truths about the world and does not advance human knowledge, as stated by Popper (103, Pg. 17): “The criterion of demarcation inherent in inductive logic—that is, the positivistic dogma of meaning—is equivalent to the requirement that all the statements of empirical science (or all ‘meaningful’ statements) must be capable of being finally decided, with respect to their truth and falsity; we shall say that they must be ‘*conclusively decidable*’. This means that their form must be such that to verify them and to falsify them must both be logically possible. Thus Schlick says: ‘. . . a genuine statement must be capable of *conclusive verification*’; and Waismann says still more clearly: ‘If there is no possible way to *determine whether a statement is true* then that statement has no meaning whatsoever. For the meaning of a statement is the method of its verification.’ It is also not a scientific theory, as Popper (103) defined it, but a pseudo-science with no meaning. This prompts us to question whether physical anthropology warrants classification as a scientific discipline. We urge the physical anthropology community to adhere to the FAIR principles, ensuring that data and methods are Findable, Accessible, Interoperable, and Reusable (FAIR), thereby facilitating thorough methodological evaluation. Ignoring FAIR principles will exacerbate the ongoing replication crisis and the public’s lack of trust in science.

The scope of our study was limited to the direct PCA usage in geometric morphometrics. However, PCA has a wide range of applications in geometric morphometrics where it is integrated into other methods, as is the case with Phylogenetic Principal Component Analysis (Phy-PCA) (44) and Phylogenetically Aligned Component Analysis (PACA) (45) or used in pathology or disciplines such as palaeontology and palaeoanthropology. Our approach can be adopted to investigate PCA accuracy in these applications.

In summary, the limitations of PCA render it ill-suitable for geometric morphometrics analyses as well as for detecting novel taxa. We showed that three ML methods (namely, nearest neighbours classifier, gaussian process classifier, and logistic regression) have superior performances for classification and highlight the effectiveness of the Local Outlier Factor (LOF) tool in identifying outlier individuals and groups in shape space. As an alternative to PCA, we created MORPHIX, an open-source Python package encompassing a comprehensive set of tools for processing superimposed landmark data. MORPHIX enables users to apply their preferred classifiers and outlier detection methods and provides a range of visualisation options. MORPHIX is freely and publicly accessible via GitHub at https://github.com/NimaMohseni/Morphometric-Analysis. We also provided detailed examples to demonstrate the usage of the package.

## Conclusions

Modern geometric morphometrics is widely used to analyse physical forms, specifically in physical anthropology. Geometric morphometrics’ heavy reliance on Principal Component Analysis (PCA) to address questions of classification as well as relationships, variations, and differences between samples and taxa, coupled with recent criticisms of this approach, merits an evaluation of its accuracy, robustness, and reproducibility. Through extensive investigations and alterations to landmark data, we showed that PCA introduces statistical artefacts specific to the data and fails to meet the required merits. We raise concerns about PCA usage in morphometrics (18,400 to 35,200 studies as of July 2024) and call for a revaluation of its usage. We demonstrate that ML methods outperform PCA for classification and outlier detection, particularly when the data are informative. We made these tools freely and publicly available in a Python package.

## Methods

### Module Description: MORPHIX

The MORPHIX Python module processes and analyses morphological data outcomes using the Morphologika software. It encompasses a range of functionalities across data processing, visualisation, and analysis. Specifically, MORPHIX facilitates the reading and processing of data saved in ’.txt’ files generated by Morphologika, extracting pertinent details such as principal component scores, eigenvalues, and percentage of total variance explained. Additionally, it offers visualisation functions employing techniques like Principal Component Analysis (PCA) plots, t-Distributed Stochastic Neighbor Embedding (t-SNE) plots, and dendrograms. Moreover, MORPHIX incorporates methods for outlier detection, classification utilising ML algorithms, and post-processing of data, thus providing a comprehensive toolkit for morphometrics analysis.

The module relies on several widely used Python libraries for data manipulation, visualisation, and ML. It offers a user-friendly interface with customisable parameters to accommodate diverse research needs. Researchers can utilise MORPHIX to analyse morphological data obtained from various sources with slight modifications. Its flexibility and scalability make it suitable for both exploratory data analysis and in-depth research investigations.

After processing landmark data in Morphologika comprising of General Procrustes Analysis and Principal Component Analysis, the output can be saved in ’.txt’ format. This module uses that text file as its input. Additionally, a data frame containing information about the samples (e.g., group, encoded labels, sex, complete sample names) is required. It is recommended to create this dataset for the initial landmark data before removing any samples, as it serves as a reference for further comparisons. The ’read’ function reads the results of a Morphologika analysis from the ’.txt’ file. The ’post_process’ function can reduce the dataset and remove certain groups if necessary. Users can also create PCA Plots and t-SNE Plots using the ’PCAplot’ and ’plot_tsne’ functions, respectively.

The example notebook in the GitHub repository demonstrates how MORPHIX can be combined with other modules, such as scikit-learn (104), for ML tasks. Researchers can leverage these integrations to perform advanced analyses and derive deeper insights from their data.

### The dataset

We used a dataset comprising the three-dimensional coordinates of 31 landmarks measured with an accuracy of 0.5 mm. The landmarks represent the locations of sutural junctions, maxima of curvature, and some other important anatomical features (Figure 15 and Table S1) of the crania of 94 adults of five papionin genera (32). Adult crania had been selected based on completed dentition (32).

The taxa were *Cercocebus* (15 specimens of *C. torquatus* including seven males and eight females), *Lophocebus* (27 specimens of *L. albigena* including 16 males and 11 females, plus two specimens of *L. aterrimus*, both female), Macaca (22 specimens *M. mulatta* including 10 males and 12 females), *Mandrillus* (20 specimens of *M. leucophaeus* including 13 males and seven females) and *Papio* (eight specimens of *P. cynocephalus* including three males and five females) (32).

The three papionins and their skull pictures presented in Figure 1 were obtained from the following links under a Creative Commons Attribution-Share Alike 3.0 Unported license with the exception of the last link, which was used with permission: https://tinyurl.com/mrh3ukz8, https://tinyurl.com/2p8777j4, https://tinyurl.com/7dxbt4zb, https://tinyurl.com/ar4uujdj, https://tinyurl.com/47n9z7ae, https://tinyurl.com/mxvr439c. All the pictures were cropped. For two skulls, the background was removed. The PC scores used for creating Figure 2 were obtained from Israel Hershkovitz. Figure 3 was obtained from Gilbert et al. (105) under the exclusive PNAS License to Publish^1^.

### Processing landmark data with General Procrustes Analysis (GPA) and Principal Component Analysis (PCA)

The process of making biological inferences from landmarks is as follows: raw landmark data consist of three-dimensional coordinates of specific points on the surface of the object or organism that are biologically meaningful or correspond to homologous structures and can be reliably and accurately located across samples. Using specialised software and hardware, the coordinates of the landmarks are recorded. The location of each landmark is recorded as a set of X, Y, (and) Z coordinates in a digital file, which can then be used for further analysis. Raw landmark data cannot be directly used as the input of computational and learning algorithms. To make the samples effectively comparable, superimposing the samples of a dataset with General Procrustes Analysis (GPA) is necessary (2, 106). This method operates in non-Euclidean shape space (107) and, through translation, rotation and scaling, superimposes the landmark data by cantering all the samples in their centroid, standardising centroid size, and reducing variations related to rotation (47). Theoretically, these steps can underline possible differences independent of position, rotation, and isometry (2). PCA is often applied after GPA to identify the major patterns of shape variation among individuals or samples. Moreover, it is used to identify the major axes of shape variation and to quantify the amount of variation explained by each PC. The first few PCs usually capture the most variation in the data and are visualised for hypothesis testing, such as comparing the shape variation between groups or observing the association between shape and other variables. The GPA and PCA analyses in this study were carried out using the morphometrics package morphologika (108).

### Employing classifiers to investigate the origin of the samples

We adopted seven ML classifiers to assess whether a new observation belongs to known taxa. Excepting for NN, this process involves a training phase in which the algorithm will try to learn the underlying patterns in the data to classify new observations. For each case, we evaluated and compared the performance of several ML algorithms, including Nearest Neighbours classifier (NN) (74), Logistic Regression (LR), Gaussian Process (GP) classifier (109), Support Vector Machine (SVM) classifier (110), Random Forest (RF) classifier (111, 112), Extra Trees (ET) classifier (113) and Extreme Gradient Boosting classifier (XGBoost) (114). We also adopted Linear Discriminant Analysis (2) for classifying the superimposed landmark data in the reduced space of the first two and 10 PCs. Tables 1 and 2 summarise their performance on the benchmark dataset and the alteration cases in terms of accuracy and balanced accuracy in a five-fold CV scheme, respectively. Supplementary Figures S1-S7 present the corresponding confusion matrices. All the post GPA-PCA analyses were carried out in python using the following modules: Numpy (115), Pandas (116), Scikit-learn (104), DEAP (117) (118) and Scipy (119). We note that while NN is not considered a traditional machine-learning method, it is commonly listed as such in the literature (120) and, for simplicity, will be referred to as such here.

### Visualisation of results

To visualise results, we used two-dimensional t-distributed Stochastic Neighbour Embedding (t-SNE) (61) for dimensionality reduction and PCA (full Procrustes projection). t-SNE aims to reduce the Kullback-Leibler divergence between the joint probability of the embedded and the high-dimensional data via translating similarities between data points into joint probabilities (61). We used two more tools in visualisation. First, convex hulls were used to define clusters of samples belonging to one taxon. Second, the decision boundary of classifiers was projected into the low-dimensional embedded space and presented with Voronoi tessellation (121) with a five-fold CV. A 1NN classifier trained on the results of the 2NN classifier was used to classify a two-dimensional mesh grid spanning the space of the plot to create the tessellation and contours used for depicting the decision boundaries.

### Kernel Density Estimation (KDE) to evaluate distributions

Kernel Density Estimation (KDE) is a non-parametric statistical method used to estimate the probability density function of a random variable. This technique involves smoothing a set of observations by creating a continuous probability density function based on a chosen kernel function (typically a bell-shaped one). The kernel function is then centred at each observation and used to estimate the probability density at that point. The estimated density is calculated by summing up the kernel functions centred at each observation point and dividing the result by the total number of observations. The bandwidth controls the smoothness of the density estimate (122). We used KDE to estimate and depict the density of the outlier scores attributed to samples by outlier detection methods in the corresponding section.

### Outlier detection methods

Outlier detection facilitates distinguishing novel samples from samples belonging to known taxa. For that, we used the Local Outlier Factor (LOF) (69) algorithm, an unsupervised method that computes the local density deviation of a given data point with respect to its neighbours. LOF considers the samples that have a substantially lower density than their neighbours as outliers. The score of the outliers (outlier) detection method was visualised with circles around samples with radii proportional to the absolute value of the scores (Eq. 1 and

Table 2). Outlier candidates were evaluated using local outlier scores (proportional to the degree to which a data point deviates from its local neighbourhood, indicating its potential as an outlier) and KDE plots. For each case, we plotted the KDE plots and a scatter plot of the t-SNE dimensions. We plotted circles around the samples, with a radius calculated according to Eq. 1 wherein *i* = 1, …, *n* is the index of the samples in the dataset, and the *score* is the outlier score according to the outlier detection method of choice.

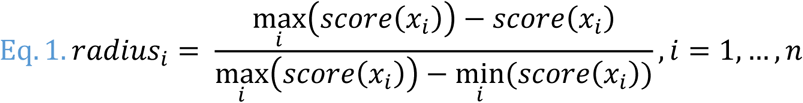

In other words, the radius around each sample is proportional to its LOF score. The final decision on whether a sample is an outlier is made by comparing the LOF scores of the sample along with other samples belonging to the potentially new taxa.

In addition to the LOF algorithm, two additional algorithms were tested. First, Isolation Forest (70), an unsupervised machine-learning algorithm used for outlier detection. It works by creating a set of isolation trees, where each tree isolates an instance in the dataset by recursively splitting the feature space. Instances that require fewer splits to be isolated are considered more anomalous, while normal instances require more splits to be isolated (70). Second, One-class SVM (71), a machine-learning algorithm used for outlier detection. It works by creating a hyperplane that separates the normal instances from the outliers in the dataset. The algorithm learns a boundary that encloses a region in the feature space where most of the normal instances lie. Instances that fall outside this boundary are considered outliers (71).

### Feature selection analysis to evaluate landmarks’ informativeness

We used a genetic algorithm as a wrapper feature selection method (77) (78) (79) (123) with a logistic regression classifier. The algorithm was implemented using the Python module ‘sklearn-genetic’ (80) based on the Python module *DEAP* (117) (118). The operator for creating the next generation in each iteration was ‘*VarAnd,’* and the type of mating between every two individuals was ‘*Uniform Crossover.’* The initial population size was set to 1000. The algorithm stops when no fitter individual is produced after five generations. We used accuracy as the objective function of the optimisation task.

### Estimating the citation number of PCA tools

Very conservatively, using Google Scholar’s citation count, we estimate that, as of July 2024, between 18,400 to 35,200 morphometric studies employed PCA without employing a conventional classification tool of the field (LDA): ‘landmarks AND morphometrics AND PCA -LDA’ (18,400), ‘morphometrics AND PCA -LDA’ (35,200). We also estimated the citation count (12,511) of PCA tools based on Google Scholar’s citation count using the following searches: ‘”geomorph” AND PCA -LDA’ (7,210) (124), ‘”morphoJ” AND PCA - LDA’ (4080) (125), ‘”TPS (Thin-Plate Spline) series” AND PCA -LDA’ (968) (126), ‘morphologika and PCA -LDA’ (590) (108), ‘”MorphpoTools” and PCA -LDA’ (153) (127), ‘EVAN Toolbox and PCA -LDA’ (132) (www.evan-society.org), respectively.

## Data and script availability

All our data and scripts that can replicate our results and figures are available via GitHub: https://github.com/NimaMohseni/Morphometric-Analysis

## Acknowledgment

EE and NM were partially funded by the Swedish Research Council award to EE (2020- 03485). The computations were enabled by resources provided by the Swedish National Infrastructure for Computing (SNIC) at Lund, partially funded by the Swedish Research Council through grant agreement no. 2018-05973. We would like to extend our heartfelt gratitude to Paul O’Higgins for sharing his landmark data and his invaluable support throughout this research. Finally, we thank the anonymous reviewers for their insights.

## Footnotes

1 Copyright (2007) National Academy of Sciences

