## Supplementary figures and tables for "Biases of Principal Component Analysis (PCA) in Physical Anthropology Studies Require a Reevaluation of Evolutionary Insights"

### The confusion matrices of the classifiers:

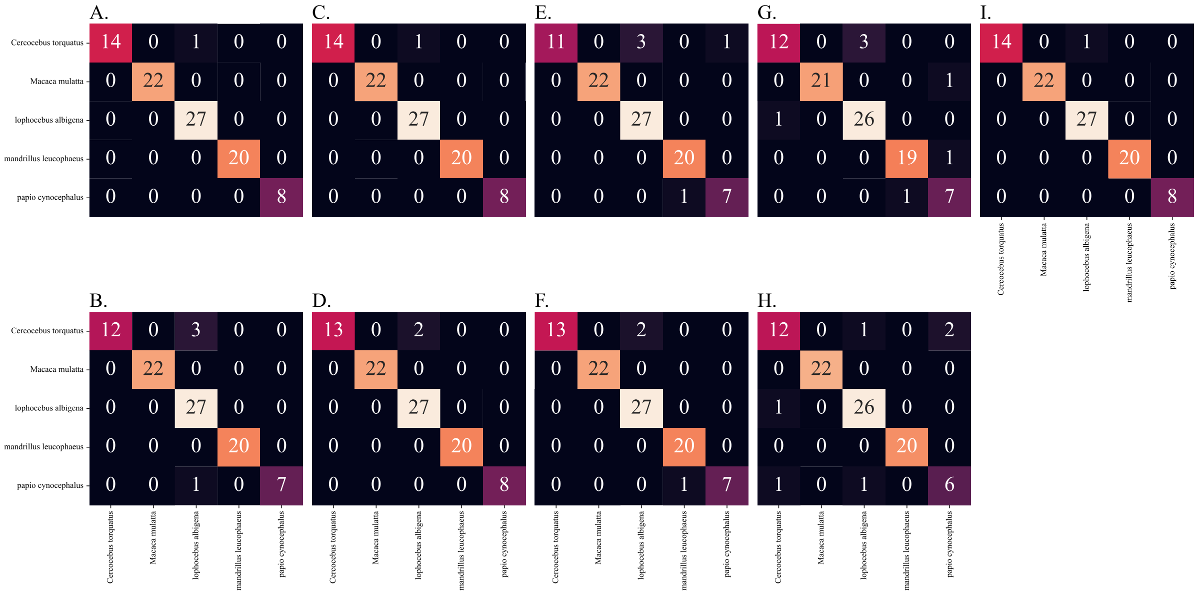


Figure S.1: Confusion matrix of the performance of the seven ML classifiers on the papionin benchmark data: A) 2NN classifier B) Logistic regression C) Gaussian process regressor D) Support vector classifier E) Random Forest classifier F) Extra trees classifier G) XGBoost classifier.

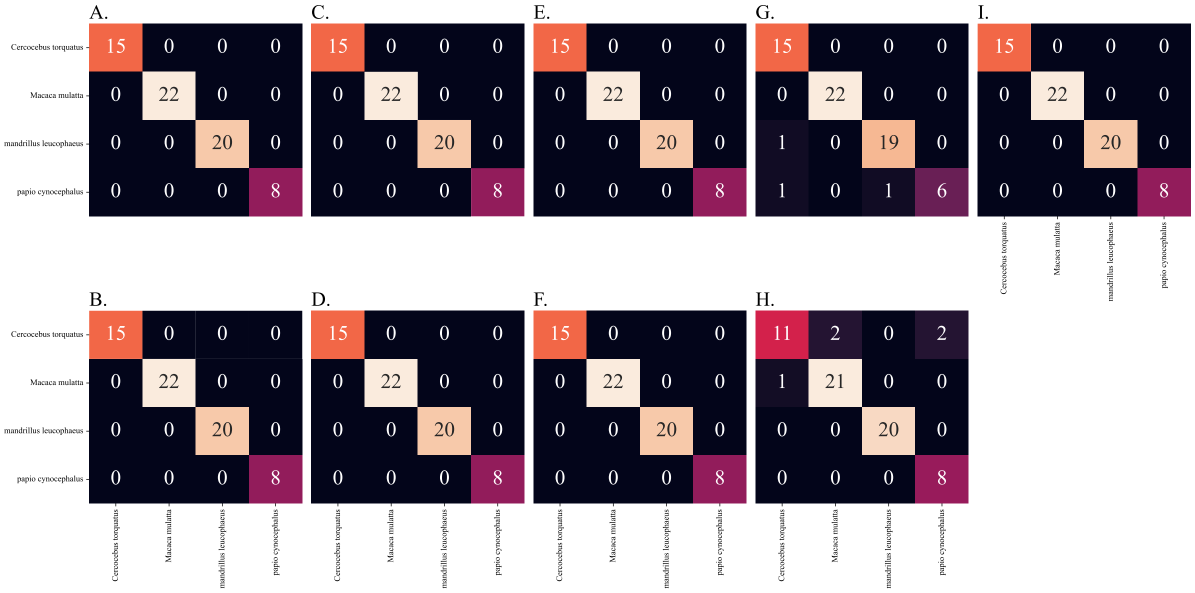

Figure S.2: Confusion matrix of the performance of the seven ML classifiers on the papionin data after the removal of the La (green) taxon. A) 2NN classifier B) Logistic regression C) Gaussian process regressor D) Support vector classifier E) Random Forest classifier F) Extra trees classifier G) XGBoost classifier.

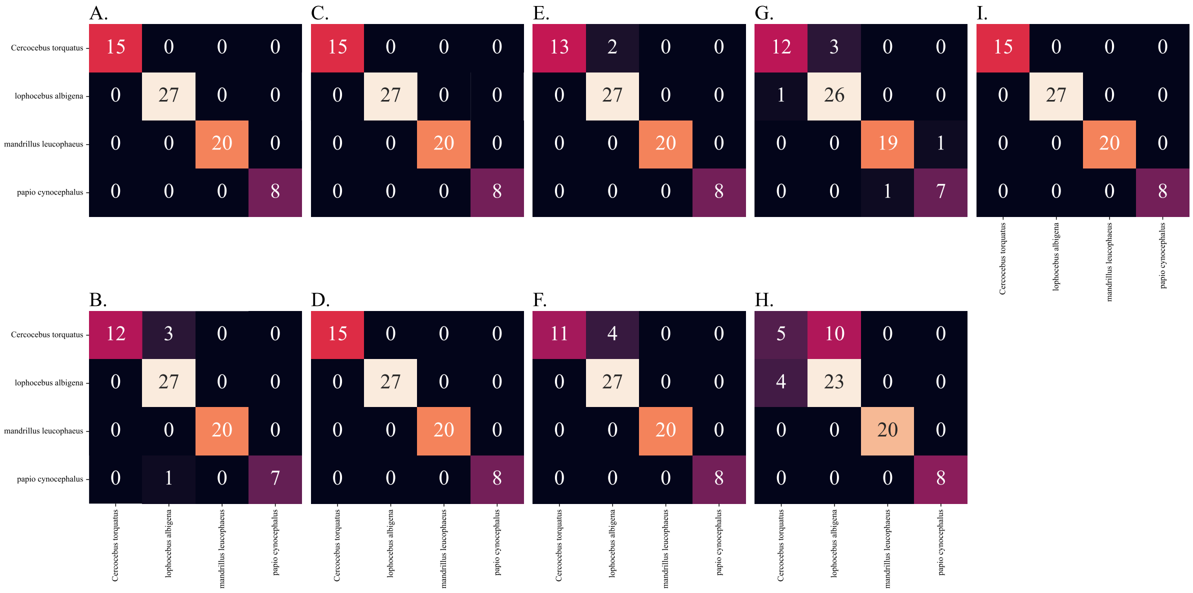


Figure S.3: Confusion matrix of the performance of the seven ML classifiers on the papionin data after removing the Mm (orange) taxon. A) 2NN classifier B) Logistic regression C) Gaussian process regressor D) Support vector classifier E) Random Forest classifier F) Extra trees classifier G) XGBoost classifier.

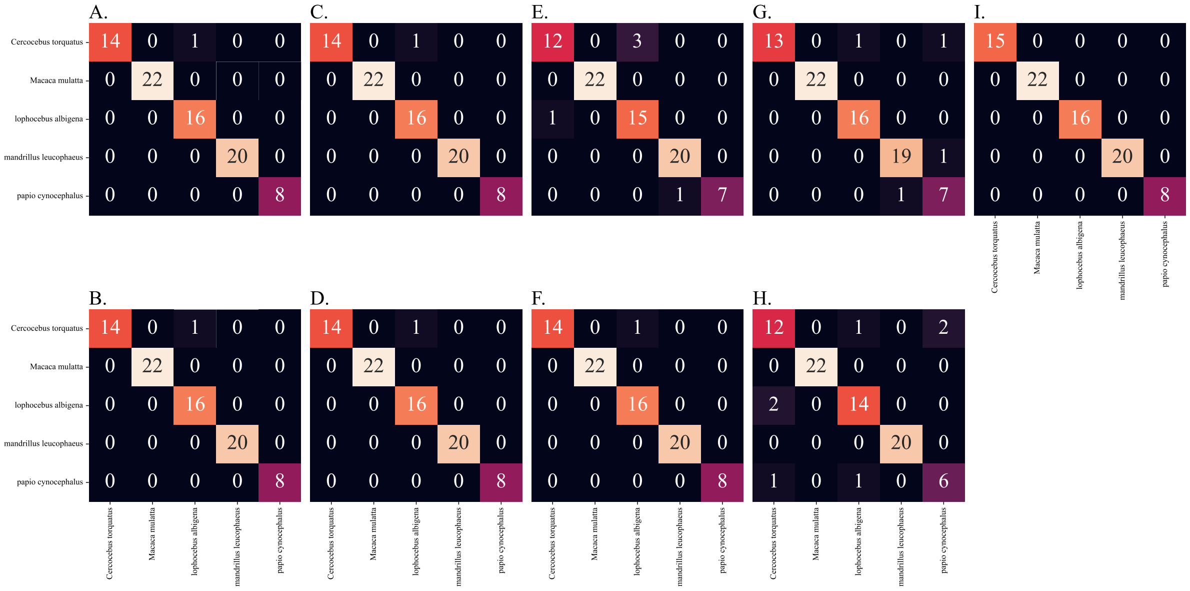

Figure S.4: Confusion matrix of the performance of the seven ML classifiers on the papionin data after the first case of sample removal. 11 *La (green)* samples (index in benchmark dataset: 52, 56, 60, 63, 64, 66, 67, 73, 75, 76 and 77). A) 2NN classifier B) Logistic regression C) Gaussian process regressor D) Support vector classifier E) Random Forest classifier F) Extra trees classifier G) XGBoost classifier.

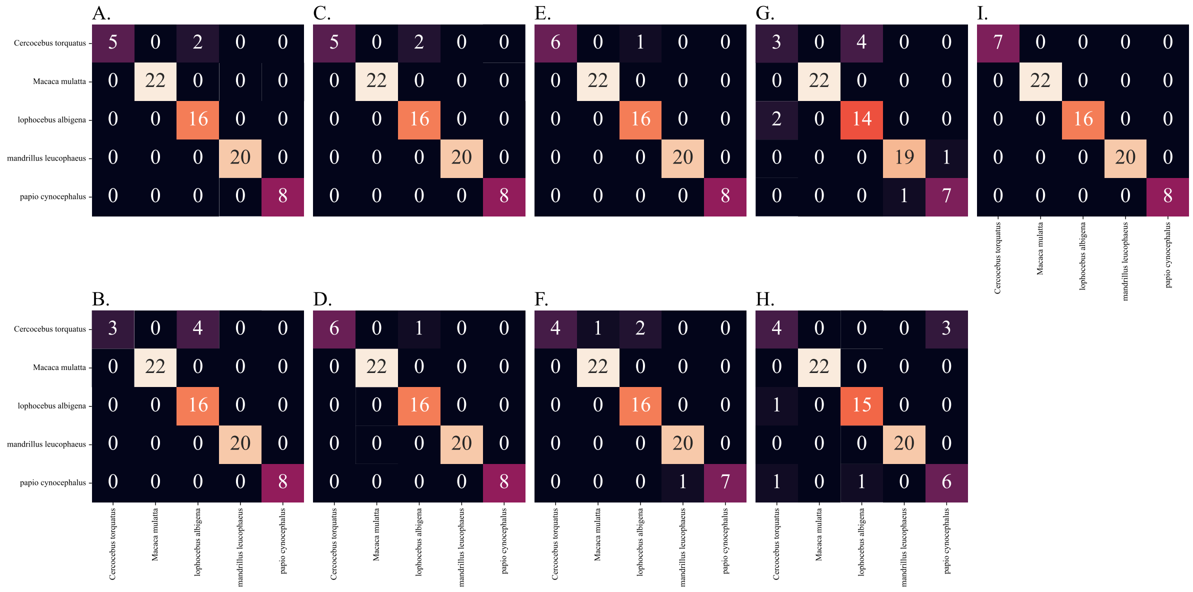


Figure S.5: Confusion matrix of the performance of the seven ML classifiers on the papionin data after the second case of sample removal. 19 samples from both *La (green)* (index in benchmark dataset: 52, 56, 60, 63, 64, 66, 67, 73, 75, 76 and 77) and *Ct (blue)* (index in benchmark dataset: 80, 81, 82, 83, 84, 88, 91 and 94) A) 2NN classifier B) Logistic regression C) Gaussian process regressor D) Support vector classifier E) Random Forest classifier F) Extra trees classifier G) XGBoost classifier.

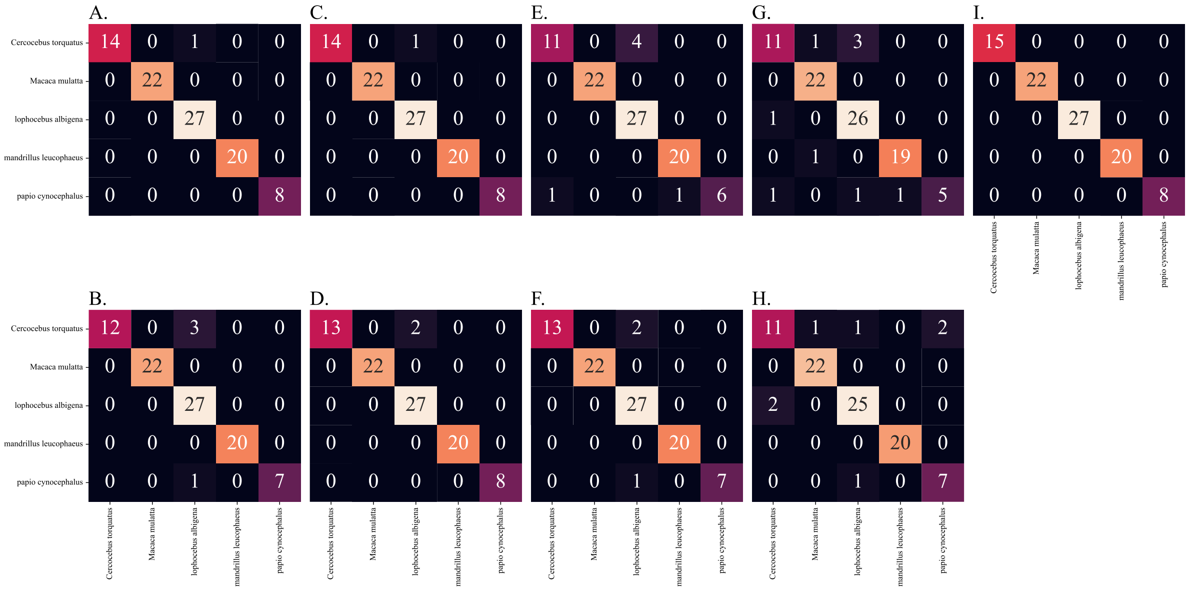

Figure S.6: Confusion matrix of the performance of the seven ML classifiers on the papionin data after the first case of landmark removal. Landmarks 23, 24, 25, 26 and 27 were removed. Scatterplots of the papionin data after the first case of sample removal. A) 2NN classifier B) Logistic regression C) Gaussian process regressor D) Support vector classifier E) Random Forest classifier F) Extra trees classifier G) XGBoost classifier.

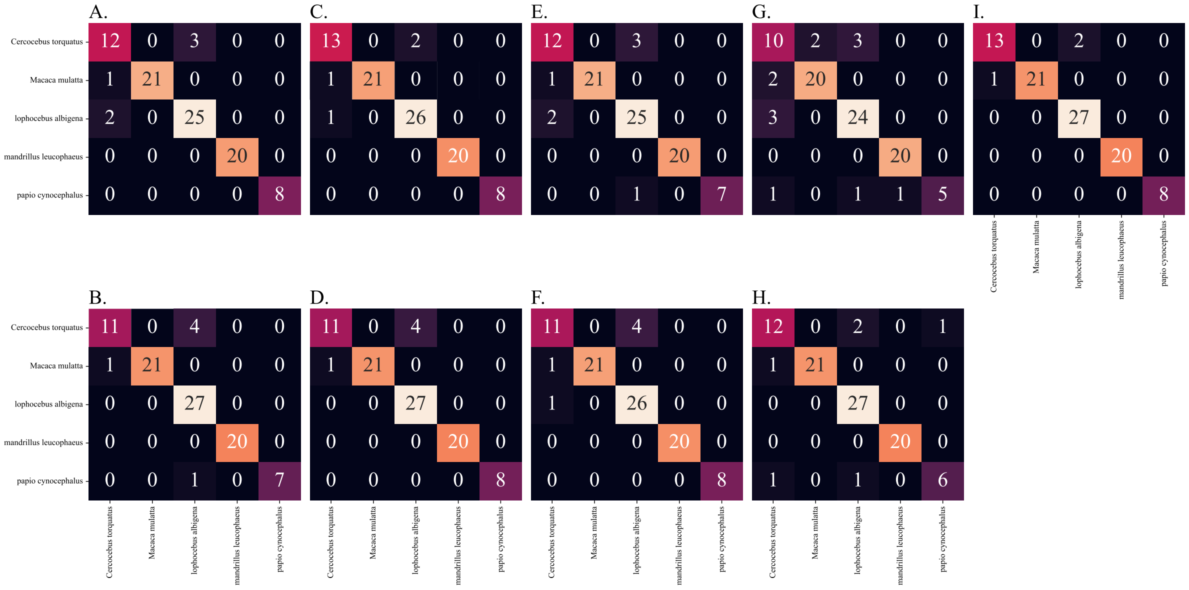

Figure S.7: Confusion matrix of the performance of the seven ML classifiers on the papionin data after the second case of landmark removal. Landmarks 11, 12, 16, 17, 18, 30 (symmetrical pair of 12) and 31 (pair of 13) were removed. Scatterplots of the papionin data after the first case of sample removal. A) 2NN classifier B) Logistic regression C)

Gaussian process regressor D) Support vector classifier E) Random Forest classifier F) Extra trees classifier G) XGBoost classifier.

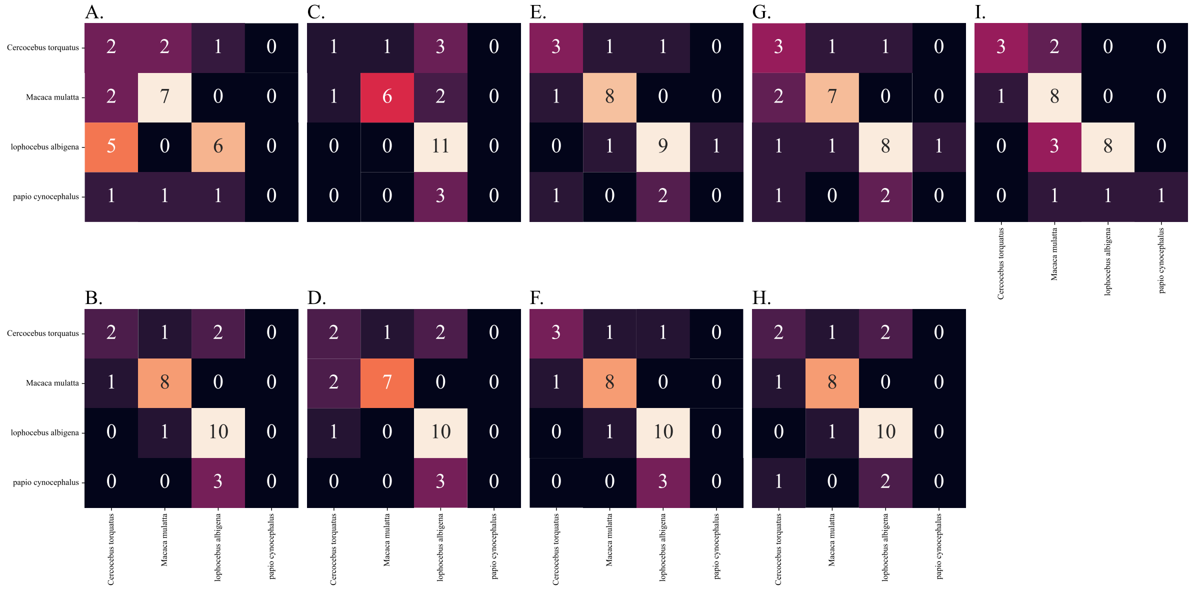


Figure S.8: Confusion matrix of the performance of the seven ML classifiers on the papionin data in the first extreme case. A) 2NN classifier B) Logistic regression C) Gaussian process regressor D) Support vector classifier E) Random Forest classifier F) Extra trees classifier G) XGBoost classifier.

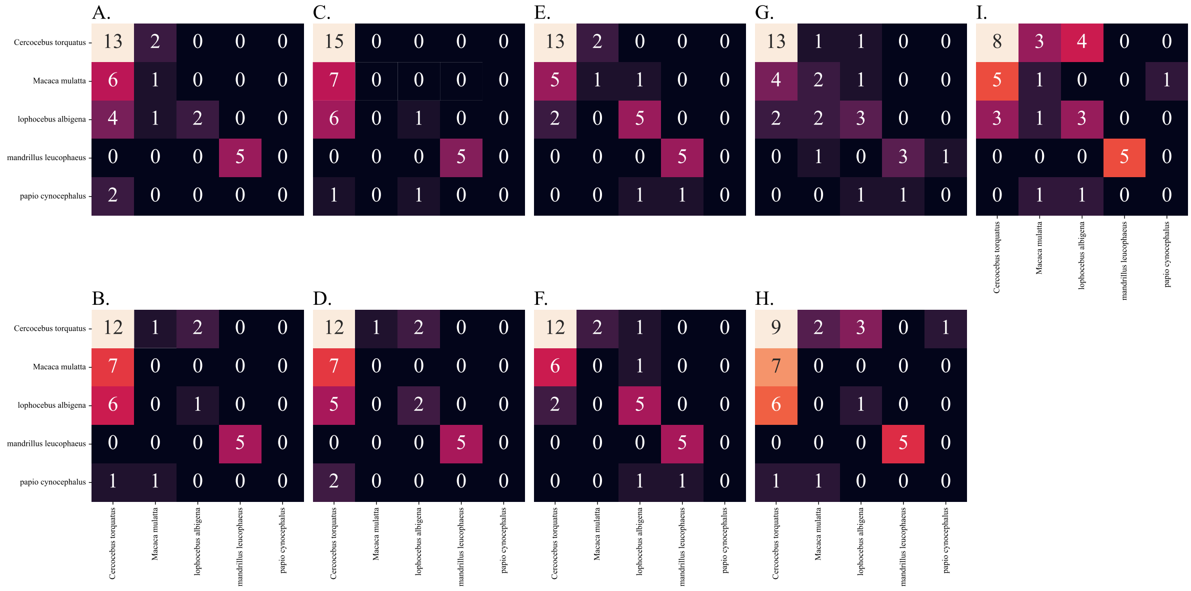


Figure S.9: Confusion matrix of the performance of the seven ML classifiers on the papionin data in the second extreme case. A) 2NN classifier B) Logistic regression C) Gaussian process regressor D) Support vector classifier E) Random Forest classifier F) Extra trees classifier G) XGBoost classifier.

#### The Scatter and KDE plots of novelty detection methods:

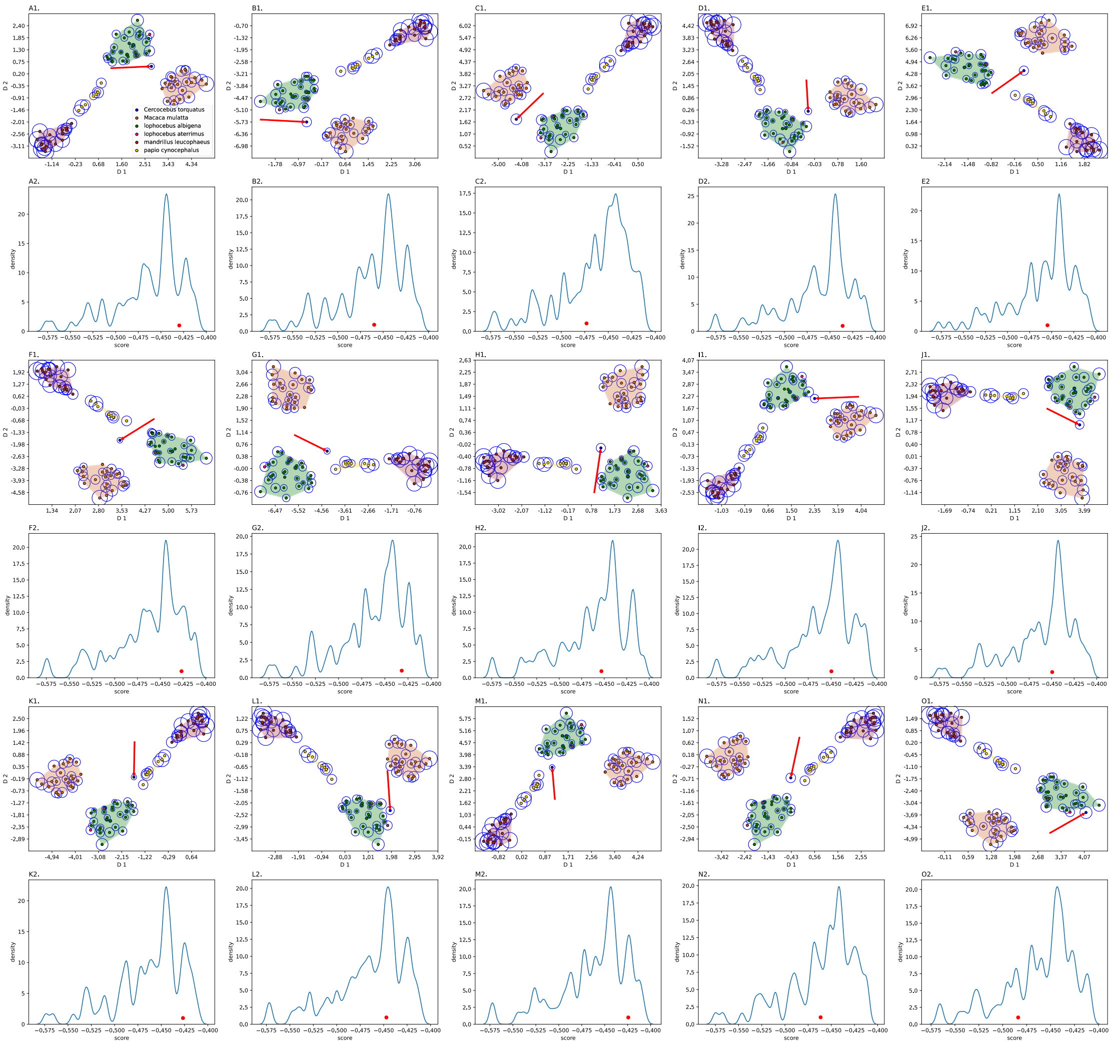


Figure S.10: Detecting outliers using isolation forest and KDE plots. Each *Ct (blue)* sample is treated as an outlier (red arrow, A1-O1) and the KDE of the scores are plotted for each case

(A2-O2). The radius of the circle around each sample is proportional to the isolation forest outlier score. The red dot in the KDE plots shows the outlier score of the outlier in each case.

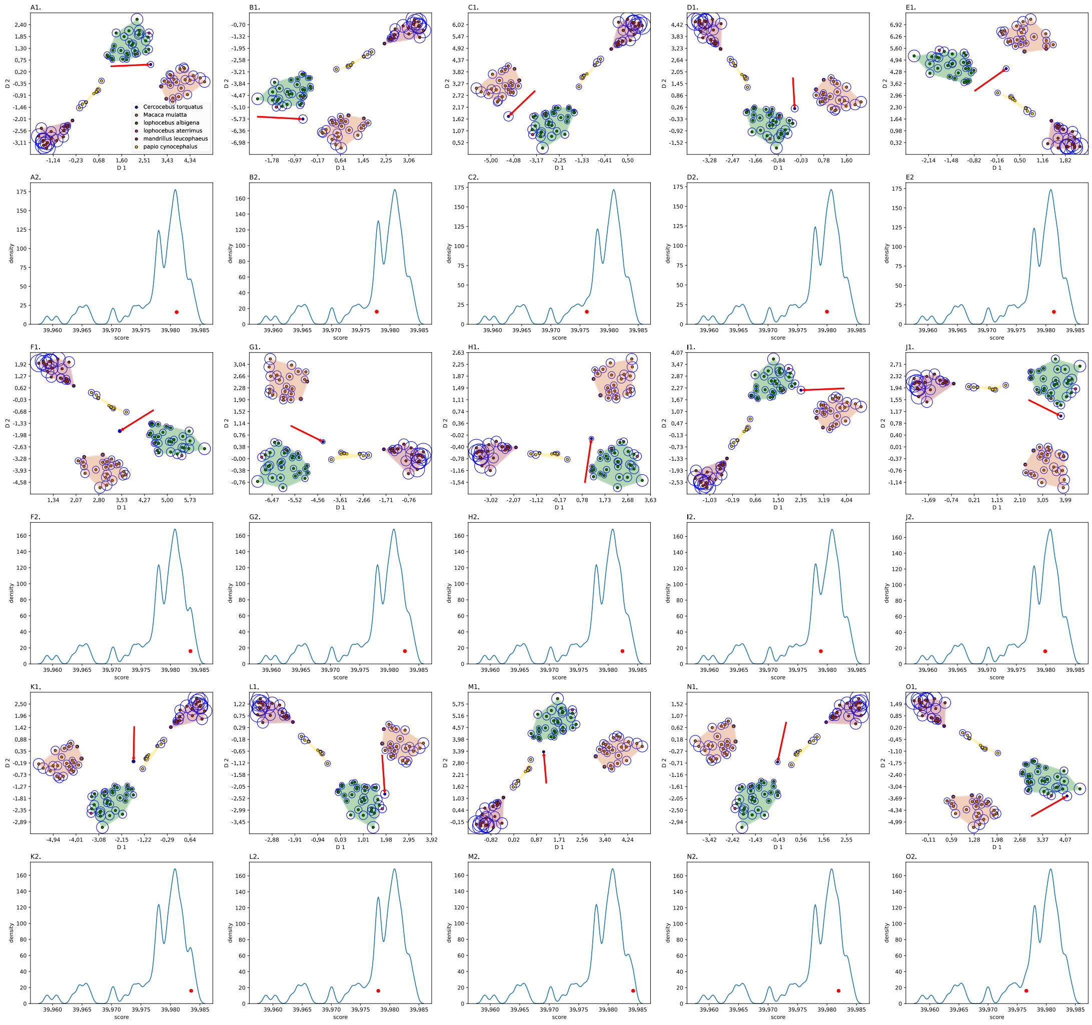


Figure S.11: Detecting outliers using one-class SVM and KDE plots. Each *Ct (blue)* sample is treated as an outlier (red arrow, A1-O1) and the KDE of the scores are plotted for each case (A2-O2). The radius of the circle around each sample is proportional to the one-class SVM outlier score. The red dot in the KDE plots shows the outlier score of the outlier in each case.

### The selected sets of landmark coordinates:

Table S1: Landmark definitions.

| Landmark number | Definition based on anatomical orientation of the face |
|---|---|
| 1 and 19 | Most lateral point on zygomatico-frontal suture on orbital rim |
| 2 and 20 | Most supero-lateral point on supraorbital rim |
| 3 and 21 | Uppermost point on orbital aperture |
| 4 and 22 | Zygomatico-frontal suture at the lateral aspect of the orbital aperture |
| 5 and 15 | Fronto-lacrimal suture at medial orbital margin |
| 6 and 24 | Zygomatico-maxillary suture at inferior orbital margin |
| 7 and 25 | Superior root of zygomatic arch |
| 8 and 26 | Inferior root of zygomatic arch |
| 9 and 27 | Zygomatico-maxillary suture at root of zygomatic arch |
| 10 and 28 | Most posterior point on maxillary alveolus |
| 11 and 29 | Deepest point in maxillary fossa |
| 12 and 30 | Maxillary ± premaxillary suture at alveolar margin |
| 13 and 31 | Nearest point to maxillary ± premaxillary suture on nasal aperture |
| 14 | Upper margin of supraorbital rim in the midline |
| 15 | Naso-frontal suture in the midline |
| 16 | Tip of nasal bones in the midline |
| 17 | Premaxillary suture at the inferior margin of the nasal aperture in the midline |
| 18 | Premaxillary suture at alveolar margin |

Table S2: The portion of informative landmarks. The second column is the accuracy of the logistic regression classifier for the selected subsets of features using genetic algorithm for each subset. The third column presents the portion of selected features.

| Subset | Accuracy | Portion of selected features |
|---|---|---|
| 1 | 0.98 | (0.41) |
| 2 | 0.98 | (0.5) |
| 3 | 0.98 | (0.43) |
| 4 | 0.98 | (0.47) |

Table S3: The selected features in five runs of the genetic algorithm.

| #Run | Selected features |
|---|---|
| 1 | False, False, False, True, False, False, False, True, False, False, False, False, True, True, True, True, False, False, True, True, False, False, True, False, False, True, False, True, False, False, False, True, False, True, False, False, True, True, False, True, False, True, True, False, False, False, True, True, False, False, False, False, True, False, True, False, False, False, True, False, False, True, True, False, False, False, True, True, False, True, True, True |
| 2 | True, True, True, False, False, False, False, True, True, True, True, True, False, False, True, False, False, False, True, True, False, True, True, True, False, True, True, False, True, False, False, False, False, True, False, False, True, False, False, False, True, True, True, False, True, True, False, False, False, True, False, True, True, True, True, True, False, False, False, True, False, False, True, True, True, False, False, False, True, False, True, False |
| 3 | False, False, False, False, False, False, True, False, False, True, True, True, False, False, True, False, False, False, True, True, False, True, True, False, True, True, False, True, False, False, False, False, False, True, True, True, False, True, False, False, False, False, True, False, |

| | |
|---|---|
| | True, False, True, True, True, True, False, False, True, False, True, False, False, False, False, False, False, False, False, False, False, False, False, True, True, False, False, True |
| 4 | False, False, False, False, True, False, False, True, True, True, False, True, True, True, True, False, True, False, True, True, False, False, True, True, True, True, True, False, False, False, False, True, False, False, True, True, False, False, False, False, True, False, True, False, True, False, False, False, False, False, False, True, True, False, True, False, False, False, False, True, True, False, False, False, False, False, True, True, True, False, False, True |
| 5 | False, False, False, True, True, True, False, True, False, True, True, False, False, False, True, True, True, False, False, False, False, True, False, False, True, True, True, False, False, False, False, True, False, True, True, True, True, True, True, False, True, False, False, False, True, False, False, True, True, False, False, False, False, False, True, False, False, False, False, False, True, True, True, False, True, True, True, True, True, False, True, False |
